# Neural bidomain model for multidimensional ephaptic coupling along general myelinated fiber bundles with nodes of Ranvier

**DOI:** 10.1101/2024.12.18.629090

**Authors:** Sehun Chun, Linyu Peng, Hae-Jeong Park

## Abstract

A novel mathematical model is proposed to investigate the distributions and functions of ephaptic coupling in generally-shaped neural fiber bundles in a multidimensional space, corresponding to the spatiotemporal interactions between propagating fiber bundles via the extracellular space. The proposed macroscopic model is derived from the classic Frankenhaeuser–Huxley model for neural spike propagation along the general neural fiber bundles in two domains (bidomain) of the nonoverlapping intracellular and extracellular space, except the common nodes of Ranvier. A high-order continuous Galerkin scheme provides an efficient two-dimensional (2D) computational simulation with moving frames (orthonormal basis vectors) representing intracellular and extracellular conductivity and nonoverlapping domains. The governing equation is mathematically and computationally validated against the existing 1D models of propagations along neural fiber bundles with the aligned nodes of Ranvier. The proposed model reveals the critical role of neural fiber bundle configurations by simulating their 2D propagations. For sufficiently large fiber bundles, synchronizing neural spike propagation along two fiber bundles with misaligned nodes of Ranvier is almost impossible. Moreover, the magnitude of interference of propagations passing-by along the fiber bundles depends on the fiber-bundle configuration, significantly depending on the location of the closest distance of two propagations, either in the myelin sheath or nodes of Ranvier. Finally, the curved fiber bundles demonstrate drastically different ephaptic coupling compared to straight fiber bundles due to the unique extracellular potential distribution by the curved fiber bundles. The new simulations highlight the importance of intrinsic fiber-bundle structures in neural spike propagation and brain functions.

**Author summary:** Fiber bundles communicate via chemical interactions across synaptic junctions and electrical interactions across gap junctions. Ephaptic coupling is another communication method between fiber bundles, which is of a small magnitude compared with other communication methods. The effect of ephaptic coupling is known on a relatively large scale and is mostly affected by the intrinsic structure of neural fiber bundles.

The study of ephaptic coupling has attracted considerable attention since the 1940s, when the term *ephaptic coupling* was first coined. However, the lack of a multidimensional model of neural spike propagation with various configurations of myelinated fiber bundles has prohibited further investigations of ephaptic coupling in brain subregions. This paper proposes a multidimensional model of neural spike simulation in the context of partial differential equations. The proposed model aims to simulate neural spike propagations in complex configurations of fiber bundles to study the functions of ephaptic coupling in such brain structures. In this paper, the new simulations of the misaligned nodes of Ranvier, interfering opposite-traveling fiber bundles, and curved fiber bundles demonstrate the critical roles of intrinsic fiber-bundle structures, including the location and alignment of the nodes of Ranvier, in neural spike propagation and brain functions.

## 1 Introduction

A multidimensional interaction of the neural spike propagation (other than chemical interactions via synaptic junctions and electric interactions via gap junctions) is achieved via the extracellular space of axons [20]. Tightly packed neural fiber bundles, the multitude of neural axons linking distributed neural populations in parallel, can laterally interact through the potential change in the extracellular space. This lateral interaction of the neural signal propagation through ion exchanges in the extracellular medium is known as *ephaptic coupling*. The neural signal transfer can be considered *digital* [47] without ephaptic coupling because the information transfer characteristics do not depend on the neural fibers pathways.

Katz and Schmitt [26] initiated the initial investigations of this phenomenon in the 1940s when they explored electric interactions between adjacent nerve fibers in *Carcinus maenas* (crabs) [33]. The term *ephaptic* was coined by Arvanitaki to distinguish it from the synaptic transmission discussed in similar research on giant axons of *Sepia officinalis* (cuttlefish) [3]. More ephaptic coupling effects (e.g., the general effect of the electric field potential in extracellular space called the*local field potential*) were observed in spinal nerve roots in dystrophic mice [42], the rat hippocampus [54], facial nerve fibers in hemifacial spasms [38], unmyelinated olfactory nerve axons [7], rat cortical pyramidal neurons [2], and the self-propagation of the longitudinal hippocampal slice [12]. Recently, the term of ephaptic coupling has been broadly employed to include more general *electric field effects* [1, 6, 9, 50] for spatiotemporal electric potential distribution in the extracellular space for neural spike propagation and the consequent neural activities.

This paper focuses on ephaptic coupling among myelinated neural fibers with nodes of Ranvier in the multidimensional extracellular space. Ephaptic coupling is mathematically and computationally modeled because analyzing the effects of the neural fiber bundle configuration for neural spike propagation is challenging in *vivo*. Neural fibers are ephaptically coupled by ions diffusing at the nodes of Ranvier, where myelin partly unwraps from the axon. With its high lipid composition, myelin insulates electricity along the axon; thus, the signal propagates along the myelinated axon with little electricity leakage. The interconnectivity of the nodes of Ranvier with the extracellular medium releases fluctuations of electric potential changes in the extracellular medium (Fig. 1). This complementary potential distribution interferes with the neural spike propagation along the neural fibers [1, 49].

**Figure 1:**
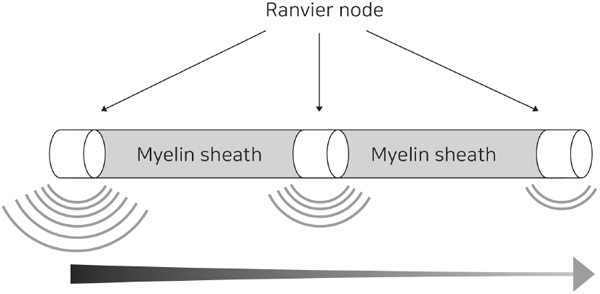
Ephaptic coupling via the neural spike propagation along a single fiber.

Mathematical models for myelinated fibers with nodes of Ranvier are often based on Frankenhaeuser and Huxley’s 1960s model of the myelinated axon [18]. This model is a later work based on the original model of an unmyelinated axon by Hodgkin and Huxley [22]. Ephaptic coupling was mathematically modeled for a three-dimensional (3D) myocardial cell in the context of cardiac electrophysiology [21, 28, 35]. Ephaptic coupling for a neural fiber deviated due to the unique structure and compositions of the neural fiber.

Goldman and Albus explored Frankenhaeuser and Huxley’s model further to demonstrate that conduction velocity is proportional to the fiber diameter [19]. In the 1970s, Waxman and Brill [8, 57, 58] and Markin [29–32] developed Frankenhaeuser and Huxley’s model further with updated parameter values to study the conduction of myelinated and unmyelinated configurations and their dynamics.

Recently, mathematical and computational models have been developed to study the interaction between myelinated fiber axon bundles with nodes of Ranvier. The conduction velocity of myelinated nerve fibers has been mathematically modeled using a simple circuit model [5]. Reutskiy et al. [43] combined a 1D model based on Frankenhaeuser and Huxley’s model with ephaptic coupling interactions to investigate the dependency of the conduction velocity on temperature and the conductivity ratio between the intracellular and extracellular space.

Schmidt and Knösche developed a mathematical framework with a spike-diffusion-spike model to confirm the effects of the configuration of the myelin sheath on ephaptic coupling [49]. The computational framework was designed to study the effect of extracellular potentials using line approximation [48] and a spike propagation model for numerous axons with various diameters [50]. Sheheitli and Jirsa applied the axonal cable theory to simulate ephaptic interactions computationally [52], and Jerez-Hanckers et al. employed a 1D cable equation for the multiscale analysis of myelinated axons [24]. A mathematical bidomain model using the FitzHugh–Nagumo dynamics was proposed for signal propagation in fascicles with many identical bundled axons [39]. All previous models were either derived in the 1D domain of a neural fiber (or fiber bundle) or could not incorporate a complex configuration of the myelin sheath and nodes of Ranvier. The previous models constrain further understanding of multidimensional ephaptic coupling in complex neural-fiber bundles.

This paper proposes a novel bidomain model simulating the multidimensional ephaptic coupling of neural spike propagation along general neural fibers. The concept of the bidomain model, first proposed by Schmitt in 1969 [51], was initially developed for neural propagation along nerve fibers [44, 55] but has become popular for propagating cardiac electric signals in the heart [41]. A multidimensional Frankenhaeuser–Huxley model is derived using the conservation of ions in the bidomain of the nonoverlapping intracellular and extracellular domains. The proposed bidomain model is unique in neural spike propagation because the two domains are nonoverlapping, and the membrane and extracellular potential should be defined separately for each domain. This model is in contrast to the bidomain model in myocardial cells where the two domains are the same. Two sets of orthogonal basis vectors, also known as *moving frames* [10, 11, 13–16], are introduced for each variable in the two domains to address this problem. The proposed multidimensional model yields the previously known mathematical and computational results in the 1D model, validating this model.

Moreover, this paper aims to provide further insight into the critical roles of fiber-bundle structures in brain functions by exploring multidimensional cases that are only possible with the proposed model. The first goal is to observe the synchronization of misaligned nodes of Ranvier, which are largely unknown because of the required multidimensional model, contrary to synchronizing myelinated fibers with the aligned nodes of Ranvier. The second goal is to observe interferences in the traveling neural spikes that are not in the same direction. Many crossing fibers in the brain demand propagational velocity changes around that region, but no analysis or simulation has been conducted. This paper simulates a simple case: oppositely traveling neural spikes in parallel fibers. The third case is the conduction velocity changes in curved fiber bundles. Brain fibers are primarily curved, and the conduction velocity is sufficiently rapid such that the curvedness of the neural fiber should affect its propagation and the corresponding synchronization. The proposed multidimensional model can simulate neural spike propagations along curved fibers to offer insight into the function of the curvedness of the fibers.

The Materials and Methods section briefly describes the 1D Frankenhaeuser–Huxley model with ephaptic coupling. Then, the novel multidimensional bidomain model is derived using the conservation of ions in the bidomain for ephaptic coupling. The numerical schemes to solve the proposed governing equations with moving frames are described with exponential error convergence results. In the Results section, the computational simulations of the neural spike propagation along a single fiber bundle and fiber bundles with aligned nodes of Ranvier confirm the previous results. Moreover, the computational results on the fiber bundles with misaligned nodes of Ranvier, opposite-traveling fiber bundles, and curved fiber bundles demonstrate the unique features of neural spike propagation. The limitations and challenges of the computational methods of the proposed model are presented in the Discussion section.

## 2 Materials and methods

### 2.1 1D Frankenhaeuser–Huxley model with ephaptic coupling

The previous mathematical and computational studies including the original model by Frankenhaeuser and Huxley [18] are primarily confined to 1D domain Ω^1*d*^ [19, 43, 49] with overlapped fibers. An axon bundle with a myelin sheath and node of Ranvier in Ω^*1d*^ is considered, where 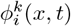 and 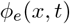 denote the electric potential of the intracellular space of the *k*th fiber and the extracellular space in Ω^1*d*^, respectively. The extracellular potential is *collective* at every point for all fibers. The membrane potential 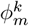 is defined as 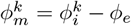. The gradient of the membrane potential proportional to the electric current is expressed as follows:

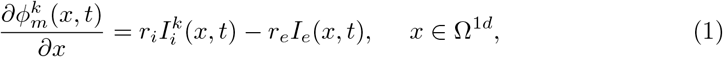

where *r*_*i*_ and *r*_*e*_ present the fiber resistance in the intercellular and extracellular media, respectively. If the values of *r*_*i*_ and *r*_*e*_ are the same for all fibers and 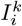 and *I*_*e*_ represent the electric current for the intercellular medium of the *k*th fiber and the extracellular medium, respectively. Substituting *I*_*e*_ = (1*/r*_*e*_)*∂ϕ*_*e*_*/∂x* into Eq. (1) yields the following intracellular electric flow for the *k*th fiber:

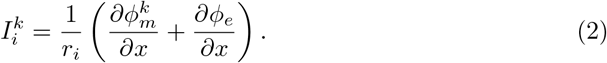

The first component is the electric flow by the gradient of the membrane potential 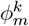. The second component is the additional current caused by the gradient of the extracellular potential *ϕ*_*e*_, known as the *ephaptic current* (point of contact) for the *k*th fiber. The electric flow of the membrane potential in the fiber is affected by the changes in the extracellular medium. In other words, two separate fibers are connected and communicated via the extracellular medium. The conservation of currents induces the dynamics of the extracellular potential *ϕ*_*e*_ contributed from all fibers as follows:

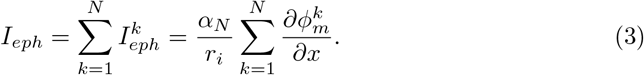

The constant *α*_*N*_ := *ρ/*(1 + *Nρ*) is called the *coupling strength constant* [43], where *ρ* denotes the conductivity ratio between the intracellular and extracellular space, defined as *ρ* = *r*_*e*_*/r*_*i*_. Moreover, *ρ* = 1 corresponds to the case in which the conductivity of the intracellular space is the same as that of the extracellular space. The coupling strength constant *α*_*N*_, which is usually less than 1.0, depends on the resistivity of the extracellular space and the surface ratio of the fiber. Eq. (3) is only possible by assuming a specific alignment between 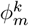 and *ϕ*_*e*_ to simplify the dynamics of ephaptic coupling in the 1D domain. Substituting Eq. (3) into Eq. (2) yields the following equation for the total current using the linearity of electric potentials:

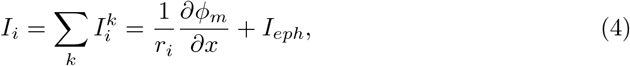

where 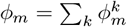. Differentiating Eq. (4) with respect to *x* and incorporating the corresponding leakage or ion channel reaction yields the following equations for the membrane potential dynamics for the myelin region:

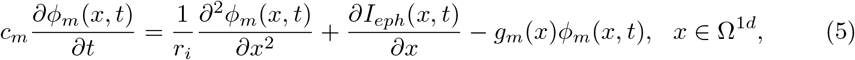

where *c*_*m*_ represents the myelin capacitance per unit length, 1.87 × 10^−11^ F/cm. In addition, *g*_*m*_ denotes the myelin conductance per unit length, 5.60 × 10^−9^ F/cm [58]. The membrane equation for the node of Ranvier is similarly derived as follows:

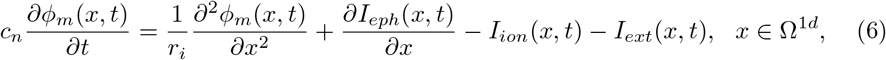

where *c*_*n*_ indicates the nodal axolemma capacitance per unit length, 3.14 × 10^−9^ F/cm, and *I*_*ext*_ represents the external current. Moreover, *I*_*ion*_ denotes the ionic current per unit length adopted from the model by Frankenhaeuser–Huxley as follows [18, 43]:

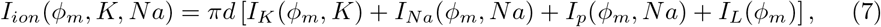

where *d* denotes the fiber diameter, *I*_*K*_ represents the potassium current, *I*_*Na*_ indicates sodium current, *I*_*p*_ denotes the non-specific delayed current, and *I*_*l*_ indicates the leak current, defined as follows:

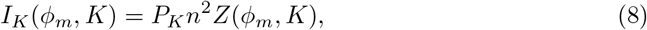

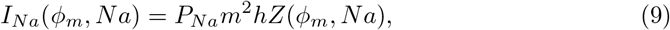

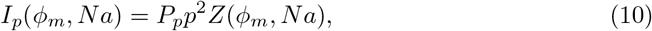

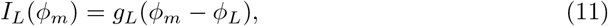

where *g*_*L*_ indicates the leak conductance, 0.0303 *mho/cm*^2^, and *ϕ*_*L*_ represents the leak current equilibrium potential, 0.026 mV. The ion variables *m, h, n*, and *p* correspond to the sodium permeability, inactivation of sodium permeability, potassium permeability, and non-specific permeability, respectively. The function *Z* in Eqs. (8) to (11) is defined as follows:

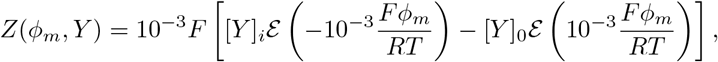

where [·]_0_ and [·]_*i*_ are the corresponding ion density values in the extracellular and intracellular space, respectively. In addition, *T* denotes the absolute temperature, *R* presents the gas constant of 8.3145 (J/mol/K), and *F* indicates the Faraday constant of 96, 484.6 (C/mol). Moreover, the function ℰ is defined as follows:

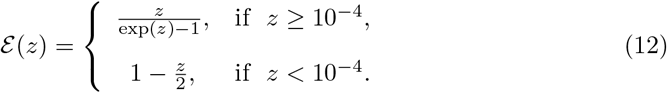

The corresponding constants and ion permeability dynamics for the variables (*m, n, p, h*) are the same as found in [18, 43, 58]. Additionally, Fig. (2) displays the time-dependent dynamics during excitation. The1D single variable model in Eqs. (5) to (6) is compact and effective for predicting and analyzing ephaptic coupling effects. However, the restriction of this model are clear. For example, Eq. (3) assumes that all nodes of Ranvier should be allocated in the exact location with the same excitation initiation along the fibers. Consequently, this 1D model is only relevant to special case studies, such as the synchronization strength of neural spikes, failing to investigate the multidimensional ephaptic coupling effects with various fiber configurations.

**Figure 2:**
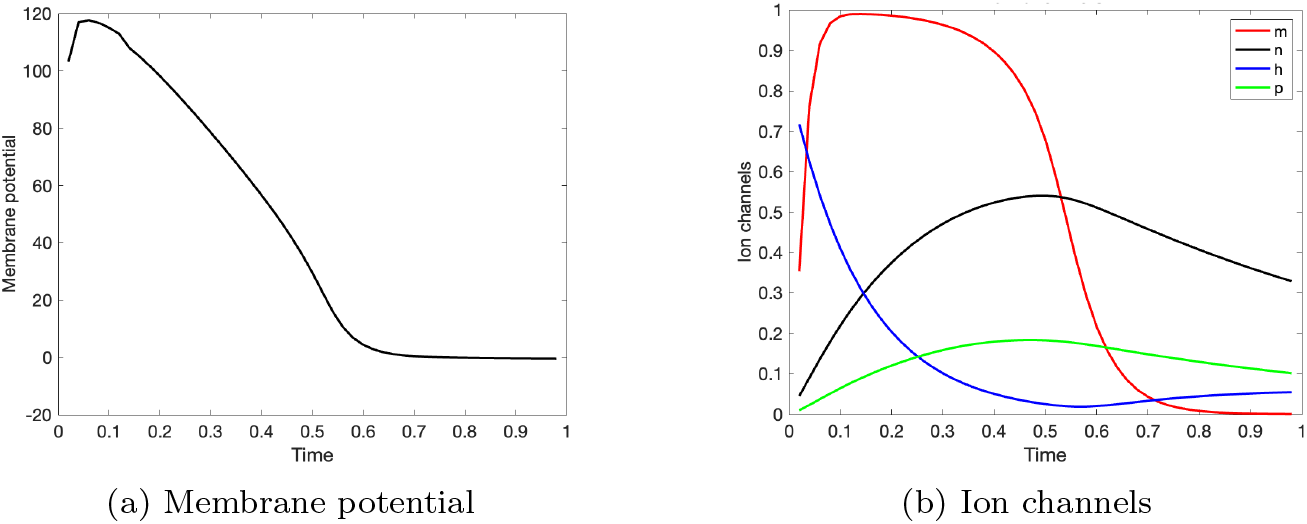
Dynamics of membrane potential and ion channels during excitation using the Frankenhaeuser–Huxely model.

### Mathematical modeling of multidimensional neural spike propagation

This work considers a two-dimensional (2D) isolated domain Ω consisting of the intracellular and extracellular space. The intracellular domain corresponds to the region neurons occupy, denoted by Ω_*i*_. The extracellular space corresponds to the space between neurons, denoted by Ω_*e*_. The intracellular domain is considered independent of the extracellular space with a clear boundary between the domains because the membrane has high resistance with thousands of Ω· cm^2^ compared to the resistance of the extracellular space with hundreds of Ω· cm^2^ [34]. The intracellular space Ω_*i*_ consists 6/33 of the nodes of Ranvier Ω_*n*_ and the internodal region Ω_*m*_ covered by a myelin sheath (Fig. 3), where

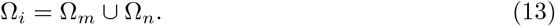

**Figure 3:**
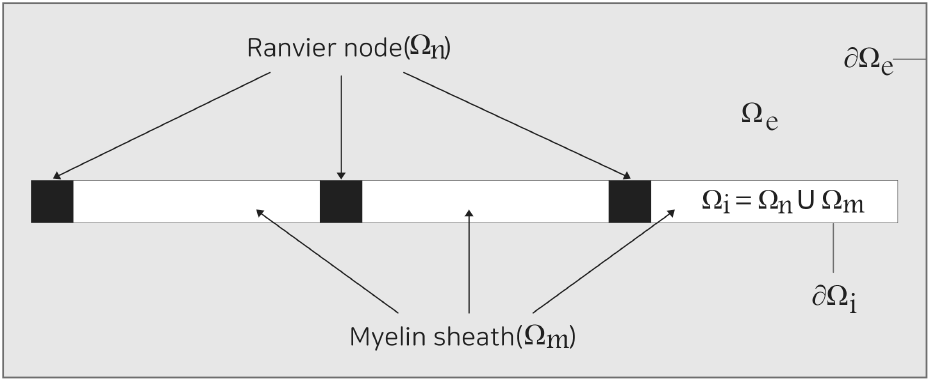
Two-dimensional fiber bundle domain Ω with the node of Ranvier Ω_*n*_ and myelin sheath Ω_*m*_. Ω_*i*_: intercellular space, Ω_*e*_: extracellular space.

Ion channel reactions primarily occur in Ω_*n*_. The extracellular space Ω_*e*_ corresponds to the area outside the intracellular domain but includes the nodes of Ranvier:

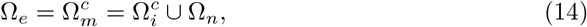

where the superscript *c* represents the complement of the corresponding domain. The intracellular and extracellular space comprises the entire domain Ω to have the node of Ranvier in common:

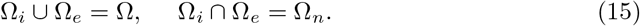

Compared to the length of the myelinated node at 2 mm, the length of the node of Ranvier is approximately 1000 times smaller about 2.5 *µ* m [43, 58]. The width of an axon is approximately 2 *µ* m, the same for the myelinated node and node of Ranvier. The boundary between the myelinated cell and node of Ranvier is not specified but is distinguished as the boundary of elements with different conductivities.

The electric current along the neural fiber is denoted by **J**_*i*_ in the intracellular region Ω_*i*_, and the conductivity tensor and electric potential are denoted by 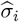 and *ϕ*_*i*_, respectively, in Ω_*i*_. The hat sign implies that the corresponding quantity is a tensor with a magnitude and direction. Note that 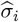 is strongly anisotropic such that a strong conductivity exists along the axon and a weak conductivity exists orthogonal to the axonal direction. Hence, the electric current along the fiber is expressed as the gradient of the electric potential *ϕ*_*i*_ along 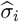 as follows:

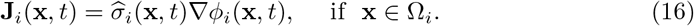

For the general 3D space, 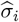 can be represented as the following matrx.

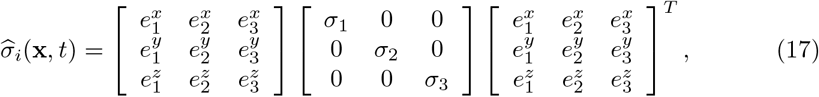

where 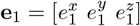 indicates the unit direction vector of the neural fiber with a conductivity of *σ*_1_. Alternatively, a curve *γ*(*s*) is considered to be the parametrization of the neural fiber with the parameter *s* within a certain range, and **e**_1_ denotes the unit tangent vector of the curve *γ*(*s*). Then, Eq. (16) is equivalently expressed as follows:

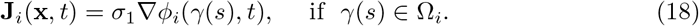

Similarly, the extracellular current **J**_*e*_ is defined in the extracellular space Ω_*e*_ with diffusivity conductivity 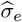 and electric potential *ϕ*_*e*_ as

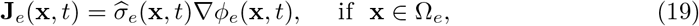

where 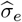 is assumed to have isotropic diffusivity with a magnitude of *σ*_*e*_. Applying the divergence operator in Eqs. (16) and (19) yields the following conservation of current densities [41]:

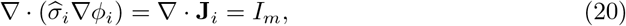

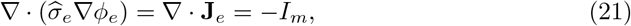

where *I*_*m*_ represents the membrane current density per unit area. The current flow across the membrane can be described by time-dependent and ionic currents as follows:

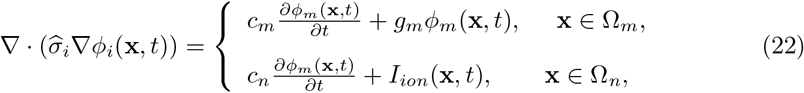

where *c*_*m*_ and *c*_*n*_ denote the corresponding capacitances, the same as those in Eqs. (5) and (6). Moreover, *g*_*m*_ indicates the myelin conductance of the myelin sheath, and *I*_*ion*_ denotes the ionic current per unit length. Substituting the definition of the membrane potential *ϕ*_*i*_ = *ϕ*_*m*_ + *ϕ*_*e*_ into Eq. (22) yields

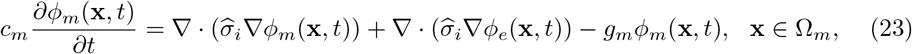

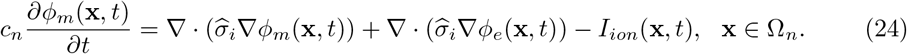

Each tessellated element is assumed to be either a myelin sheath or the node of Ranvier, not both regions in a single element. Table 1 summarizes each region, defining each variable and tensor.

**Table 1:** Regions defining each variable and tensor (*O*: defined: *X*: not defined).

| | $\phi_i$ | $\phi_e$ | $\phi_m$ | $\hat{\sigma}_i$ | $\hat{\sigma}_e$ |
| --- | --- | --- | --- | --- | --- |
| Node of Ranvier | O | O | O | O | O |
| Myeline | O | O | O | O | X |
| Extracellular space | X | O | X | X | O |

To derive the conservation of ions in Ω, each variable *ϕ*_*i*_ ∈ Ω_*i*_ and *ϕ*_*e*_ ∈ Ω_*e*_ defined in each non-overlapping domain is expanded into 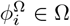 and 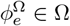 in the entire domain Ω as follows:

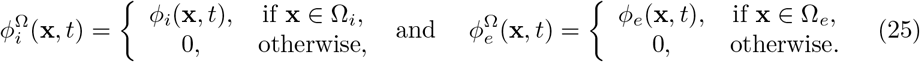

Similarly, the conductivity tensor of 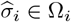 and 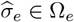 are expanded into 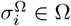 and 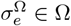 as

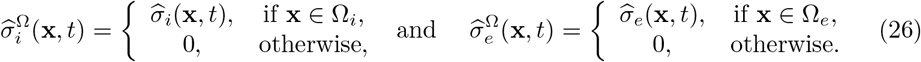

Adding Eqs. (16) and (19) with Eqs. (25) and (26) yields the following equation for the conservation of ions:

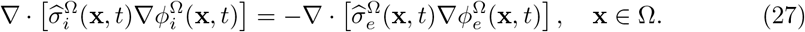

For the electric potential *ϕ*, the corresponding quantity of 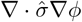 represents the current source density corresponding to the overall charged ion density [34, 37, 40]. The left- and right-hand sides of Eq. (27) corresponds to the current source density of the intracellular and extracellular space, respectively. Thus, Eq. (27) states that the sum of the current source density of the intracellular and extracellular space is zero, equivalent to the conservation of ions in a domain with two separate spaces. Substituting *ϕ*_*i*_ = *ϕ*_*m*_ + *ϕ*_*e*_ into Eq. (27) results in the following equations.

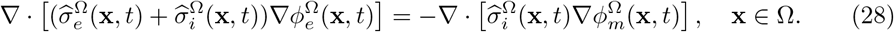

Equations similar to Eqs. (23), (24) and (28) are known as *bidomain equations*, particularly in cardiac electric signal propagation [25, 41, 45]. The 2D decompositions of tensors 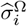 and 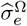 are obtained as follows, assuming a cylindrical shape of the fiber with a circular base of radius *a*:

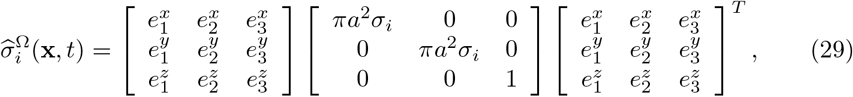

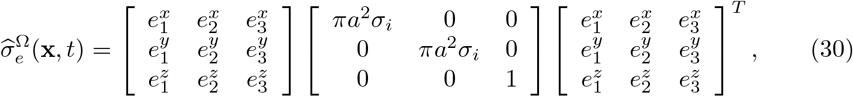

where *σ*_*i*_ and *σ*_*e*_ denote the maximum conductivity in the intracellular and extracellular space, respectively. Substituting Eqs. (29) and (30) into Eq. (28) results in the following Poisson equation along the fiber direction **e**_1_ and its orthogonal direction **e**_2_ for **x** ∈ Ω,s

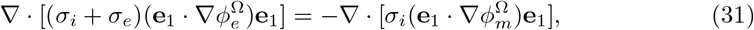

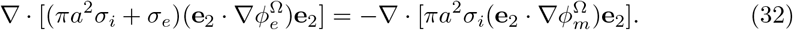

Moreover, another variable is introduced to approximate the case with a significant extracellular potential, compared to the membrane potential, without including more fibers. If the strength of the extracellular potential is proportional to the square of the fiber bundle radius, *r*_*b*_, as modeled in [48], which is continuously filled with axons of radius *a*, then the cumulative extracellular potential at *x* outside the fiber can be derived as follows:

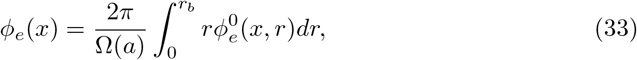

where Ω(*a*) is the cross-sectional area filled by the axons of radius *a* and 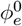 represent the magnitude of the extracellular potential of each fiber. If each induced extracellular potential by every axon is constant as 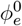 along the radial direction, then the following expression defines the extracellular potential inside the fiber bundle.

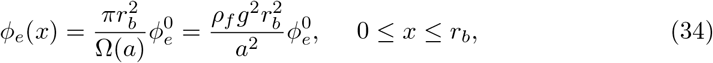

where Ω(*a*) = *πa*^2^*/*(*ρg*^2^) is applied for the constant *ρ*_*f*_ as the relative size of the volume occupied by the fibers, and *g* is the ratio between the axon diameter and fiber diameter (the *g*-ratio). Substituting Eq. (34) in Eq. (32) results in the multidimensional Poisson equation:

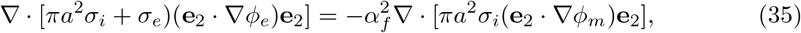

where *α*_*f*_ denotes the amplified factor of the extracellular potential, defined as 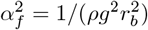 [48].The increased fiber bundle radius (*r*_*b*_) does not increase the conduction velocity by increasing the threshold of ion channels. The parameter *d* corresponding to the fiber diameter in Eq. (7) is constant, regardless of *r*_*b*_, in the reaction functions *I*_*ion*_ at the nodes of Ranvier. However, the slight change in conduction velocity by various fiber radii is indirectly caused by the change in the extracellular potential and induced electric currents.

#### Validation of the governing equations

This work considers the dynamics of the membrane potential propagation, which is affected by the distribution of the extracellular potential of the node of Ranvier, to validate the bidomain equations (Eqs. (23) and (24)).

##### Proposition 1.

*The bidomain equations (Eqs*. (23) *and* (24)*) along the fiber direction at the node of Ranvier in the 1D domain are equivalent to the 1D ephaptic coupling equations, as presented in Eqs*. (5) *and* (6) *with Eq*. (3).

*Proof*. The 1D projections of the multidimensional governing equation in Eqs. (23) and (24) are given for the 1D domain Ω^1*d*^ as follows:

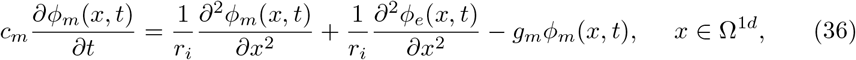

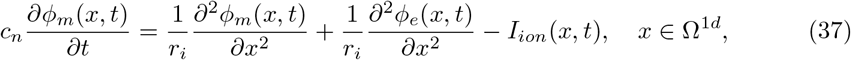

where *σ*_*i*_ = 1*/r*_*i*_. With the conductivity ratio *ρ* such that *ρ* = *σ*_*i*_*/σ*_*e*_, Eq.(28) along the fiber at the node of Ranvier is expressed as follows:

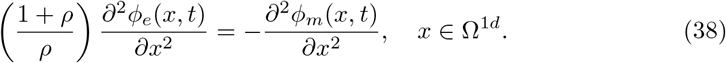

Substituting Eq. (38) into Eqs. (36) and (37) results in

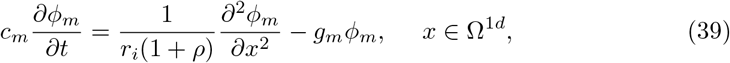

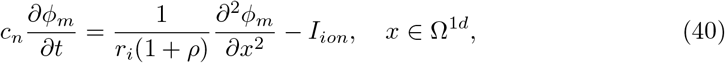

which are the same as Eqs. (5) and (6) using Eq. (3).

Moreover, the dynamics orthogonal to the fiber direction at the node of Ranvier are considered, as indicated in Eq. (32).

##### Proposition 2.

*The extracellular potential of Eq. (32) parallel to* **e**_2_, *or orthogonal to the fiber direction* **e**_1_ *at the node of Ranvier, is equivalent to the extracellular potential for a sufficiently small axon radius a as follows [48]:*

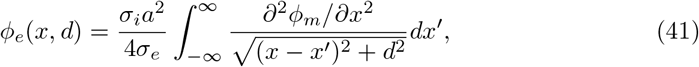

*where x denotesss the axis whose tangent vector is the same as* **e**_2_. *Moreover, d represents the distance from the axon, and a indicates the axon radius*.

*Proof*. For a constant *δ*, the solution to the scalar potential of ∇^2^*ϕ*_*e*_ = −*δ* along an infinite line is derived as follows [23, 48]:

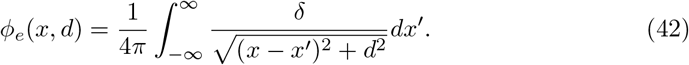

The electric potential is obtained by setting *d* → 0. Using Eq. (42) for Eq. (32) yields the following expression for *δ*,

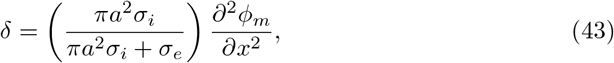

where **e**_2_·∇*ϕ*_*m*_ = *∂ϕ*_*m*_*/∂x*. If *σ*_*i*_ and *σ*_*e*_ are approximately constant in each fiber, the solution of Eq. (32) is derived as follows:

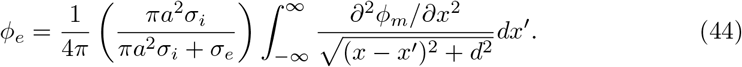

For a significantly small fiber bundle radius (*r*_*b*_), to obtain Eq. (41), *a* ≪ 1.0 and *πa*^2^*σ*_*i*_ + *σ*_*e*_ ≈ *σ*_*e*_.

### Numerical schemes with moving frames

A set of orthonormal basis vectors (**e**_1_, **e**_2_, **e**_3_) in three-dimensional (3D) space, called a *moving frame* [10, 11, 13–16], is employed for the efficient numerical solution of the bidomain equations in Eqs. (23) to (28). A moving frame can be considered a hexahedron at point *P* with each edge aligned along the unit tangent vectors (Fig. 4). A moving frame is denoted by {**e**_1_, **e**_2_, **e**_3_} satisfying the following orthonormality.

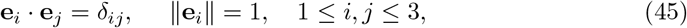

where *δ*_*ij*_ denotes the Kronecker delta. Two tangent vectors **e**_1_ and **e**_2_ lie on the tangent plane of a 2D domain Ω, whereas **e**_3_ is aligned along its surface normal vector,.

**Figure 4:**
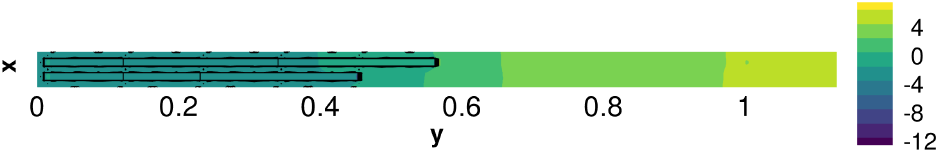
Orthonormal (orthogonal with unit length) basis vectors, called moving frames.

By introducing the capacitance ratio 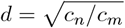, Eqs. (23) and (24) change to

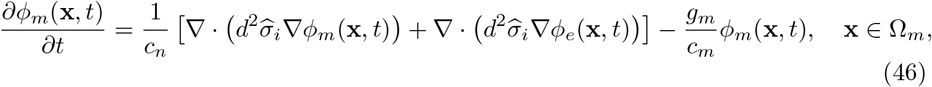

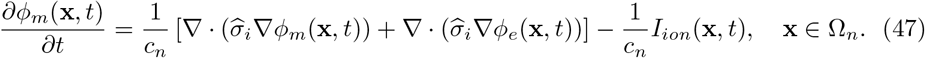

The following lemma is considered using the rescaled moving frames.

#### Lemma 1.

*In a 2D domain* Ω, *the diffusion operator with a conductivity tensor* 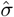 *is equivalently computed by rescaling the moving frames as follows:*

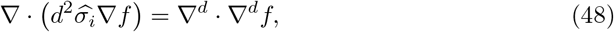

*where* ∇^*d*^ *represents the differential operator with orthogonal basis vectors* {**d**_*j*_}*of a magnitude of d, defined such that*

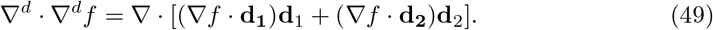

*Proof*. If orthogonal basis vectors {**e**_1_, **e**_2_} correspond to the diffusivity tensor 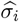, then 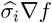 can be computed using the orthogonal basis vectors {**e**_1_, **e**_2_} as follows:

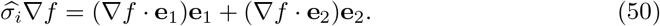

Similarly, the left-hand side of Eq. (48) can be computed as follows:

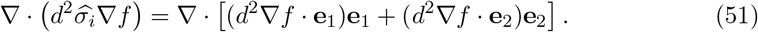

A scaled basis vector **d**_*j*_ is introduced such that

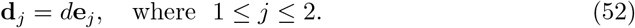

Then, Eq. (51) becomes

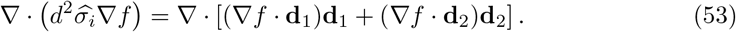

The right-hand side of Eq. (53) is the expression of the diffusion operator with the orthogonal basis vectors {**d**_*j*_}, which is the same as the right-hand side of Eq. (48).

By Lemma 1, the diffusion-reaction equations in Eqs. (46) and (47) and the anisotropic Poisson equation in Eq. (28) yield the following equations, called *neural bidomain equation*:

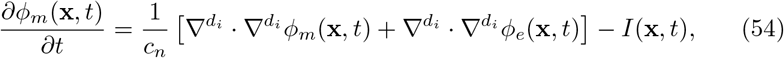

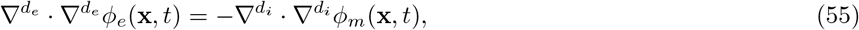

where *d*_*i*_ and *d*_*e*_ represent the magnitude of the corresponding tangent vectors {**e**_*j*_} for the intracellular and extracellular conductivity tensors 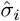 and 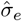, respectively, as presented in Table 2 and Fig. 5 for each domain.

**Table 2:**
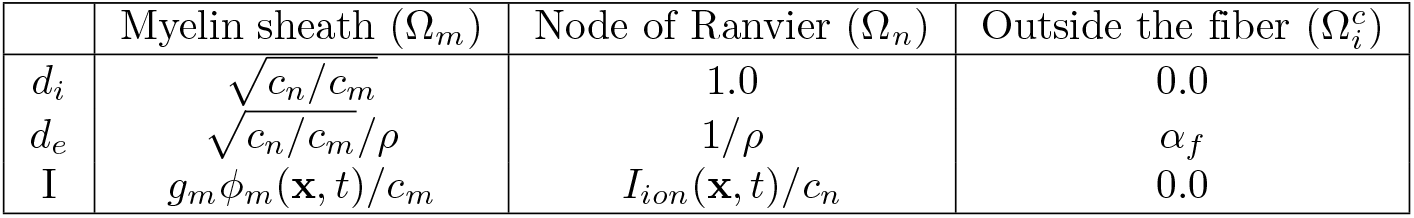
Magnitude of the tangent vectors of the moving frames for intracellular and extracellular conductivity tensorss in each domain and the corresponding electric currents (*I*). In addition, *d*_*i*_ and *d*_*e*_ corresponds to the intracellular and extracellular conductivity tensors 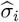 and 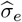, respectively. Moreover, *ρ* represents the conductivity ratio, and *α*_*f*_ denotes the amplified factor of the extracellular potential.

| | Myelin sheath ($\Omega_m$) | Node of Ranvier ($\Omega_n$) | Outside the fiber ($\Omega_i^c$) |
| --- | --- | --- | --- |
| $d_i$ | $\sqrt{c_n/c_m}$ | 1.0 | 0.0 |
| $d_e$ | $\sqrt{c_n/c_m}/\rho$ | $1/\rho$ | $\alpha_f$ |
| I | $g_m \phi_m(\mathbf{x}, t)/c_m$ | $I_{ion}(\mathbf{x}, t)/c_n$ | 0.0 |

**Figure 5:**
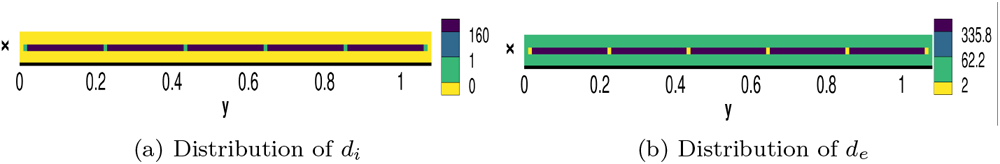
Distribution of the magnitude of the tangent vectors of *d*_*i*_ and *d*_*e*_ corresponding to the intracellular and extracellular conductivity tensors 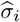 and 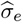, respectively, for a single fiber with *ρ* = 1.

#### Weak formulation

The following weak formulation is applied in the context of the Galerkin formulation to solve the partial differential equations of Eqs. (54) and (55). For a given test function φ of the same polynomial order as the solution, the following Galerkin formulation is computed for Eq. (54):

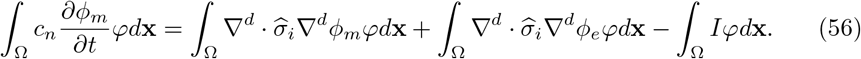

The boundary conditions are considered as follows. The intercellular space is distinguished from the extracellular space by the membrane, represented by the boundary *∂*Ω_*i*_. The electric insulation property of the membrane is expressed by the pure Neumann boundary condition as

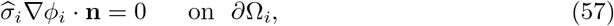

where **n** indicates the edge normal vector of the elements orthogonal to the boundary. The following equality, *ϕ*_*i*_ = *ϕ*_*m*_ + *ϕ*_*e*_, obtains the boundary condition of Eq. (57) on *∂*Ω_*i*_:

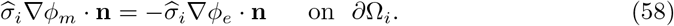

If the conductivity tensor of the neural fiber is significantly larger in the orthogonal direction to **n** and the fiber is sufficiently thin compared to its length, then, the external potential change in the anisotropic fiber along the normal direction is almost negligible:

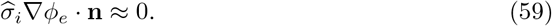

Therefore, the boundary condition in Eq. (58) is approximated as follows:

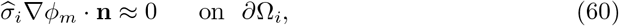

Moreover, this work considers the boundary conditions of the extracellular space Ω_*e*_, or the space surrounding the cells. The outskirt boundary of the domain Ω_*e*_ is also insulated from any influx to have the following boundary condition:

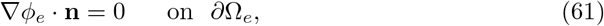

where the extracellular space is assumed to be isotropic. Furthermore, the following numerical schemes are introduced to address the numerical challenges, particularly regarding *numerical stiffness*. One difficulty in computationally solving Eq. (56) is the dramatic ratio between the node and myelin sheath length. The internodal length is approximately 2,000 *µ* m when the nodal length is 2.5 ∼3.183 *µ* m [58] [43]. Thus, the internodal region and node ratio is approximately between 1/200 and 1/1000. A minimal node sizeΔ*x* is required, affecting the time-step size for the numerical solution of the corresponding partial differential equations. The time-step size iin the context of the finite difference scheme is approximately proportional to Δ*x*^2^ [27, 56]. A high-order implicit-explicit time marching scheme is adopted to alleviate the strong restriction of the time step [4]. Eq. (56) becomes the following by considering the equation at each time.

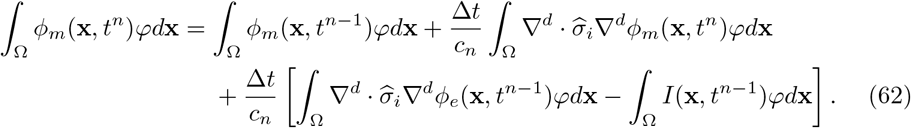

The Laplacian operator for *ϕ*_*m*_ should be solved *implicitly* to be computed at time *t*^*n*^, leading to a large global matrix that must be solved iteratively for a given time *t*^*n*^. This implicit scheme allows an approximately 10 ∼ 100 times larger time step than the explicit Laplacian operator for *ϕ*_*m*_. The other components, such as the Laplacian operator for *ϕ*_*e*_, the reaction function, and external flow **I**, are time-marched *explicitly*. The explicit Laplacian operator for *ϕ*_*e*_ affects the time step less significantly than that for *ϕ*_*m*_, possibly due to the smooth distribution of *ϕ*_*e*_.

Another stiffness problem arises for the integration of the reaction function. For the four variables for permeabilities (*m, n, p, h*), we have the following:

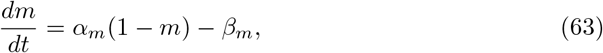

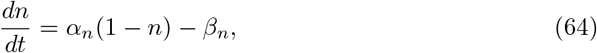

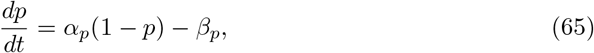

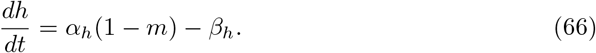

As illustrated in Fig. 2, the dynamics of ion channels demonstrate sharp variations with respect to membrane potential changes. This stiffness in the reaction functions also significantly restricts the time-step size. The Rush–Larsen scheme [46] was adopted for the numerical scheme to alleviate the strong restriction by the stiff reaction function. Applying the Rush–Larsen scheme to Eqs. (63) - (66), the solution at the *i*th time stage for *A* = (*m, h, p, h*) is obtained as follows:

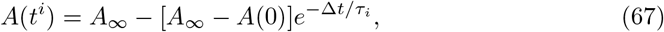

where *A*_*∞*_ = *α*_*A*_*/*(*α*_*A*_ + *β*_*A*_) and *τ*_*i*_ = 1*/*(*α*_*A*_ + *α*_*B*_).

#### Anisotropic Helmholtz Solver

In the numerical solution of Eqs. (54) and (54), the following 2D Helmholtz equation is solved repeatedly, for example, in the implicit time marching of Eq. (54) and the anisotropic Poisson equation for the extracellular potential in Eqs. (31) and (32):

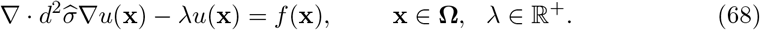

This work considers one tessellated element Ω^*k*^ of a 2D2 domain Ω such that ∩_*k*_Ω^*k*^ = Ω and ∪_*k*_Ω^*k*^ = ∅. For each Ω^*k*^, Eq. (68) is integrated by part after multiplying a test function φ of the same polynomial degree as the solution function *u*, obtaining

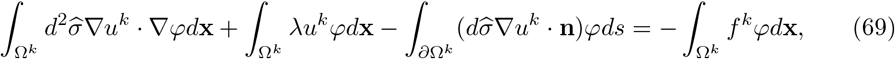

where *u*^*k*^ and *f* ^*k*^ are the projections of *u* and *k* on the element Ω^*k*^, respectively. An alternative expression is obtained by Lemma 1 as follows:

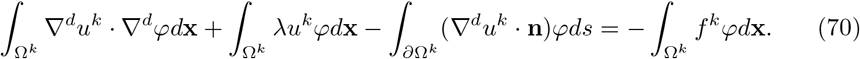

An axis 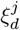 is defined such that it satisfies the following expression of directional derivative along **d**_*j*_, that is,

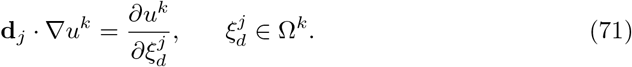

Using Lemma 1 again to incorporate the diffusivity tensor into moving frames yields the following discretization of the anisotropic Helmholtz solver for the solution *u*^*k*^ in an element Ω^*k*^:

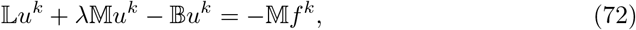

wher

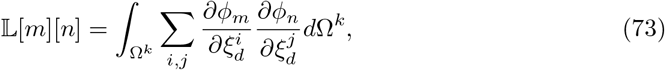

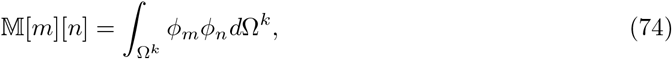

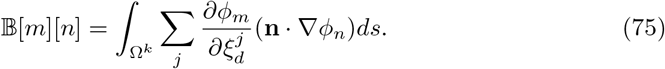

The following test problem is employed to demonstrate the accuracy of the proposed anisotropic Helmholtz solver in Eq. (72) in a 2D domain of (*x, y*) ∈ [−1, 1] × [−1, 1].

The following Helmholtz equation with an tensor 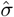 is considered:

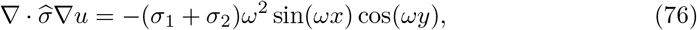

where the conductivity tensor is given by

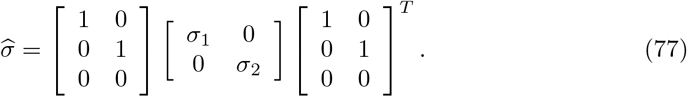

The exact solution of this test problem is *u* = sin(*ωx*) cos(*ωy*). In addition, Fig. 6 displays the exact solution with *ω* = *π* and the exponential error convergence for the isotropic case with unit conductivity (*σ*_1_ = *σ*_2_ = 1.0), the isotropic case with nonunit conductivity (*σ*_1_ = *σ*_2_ = 4.0), and the anisotropic case (*σ*_1_ = 0.5, *σ*_2_ = 4.0).

**Figure 6:**
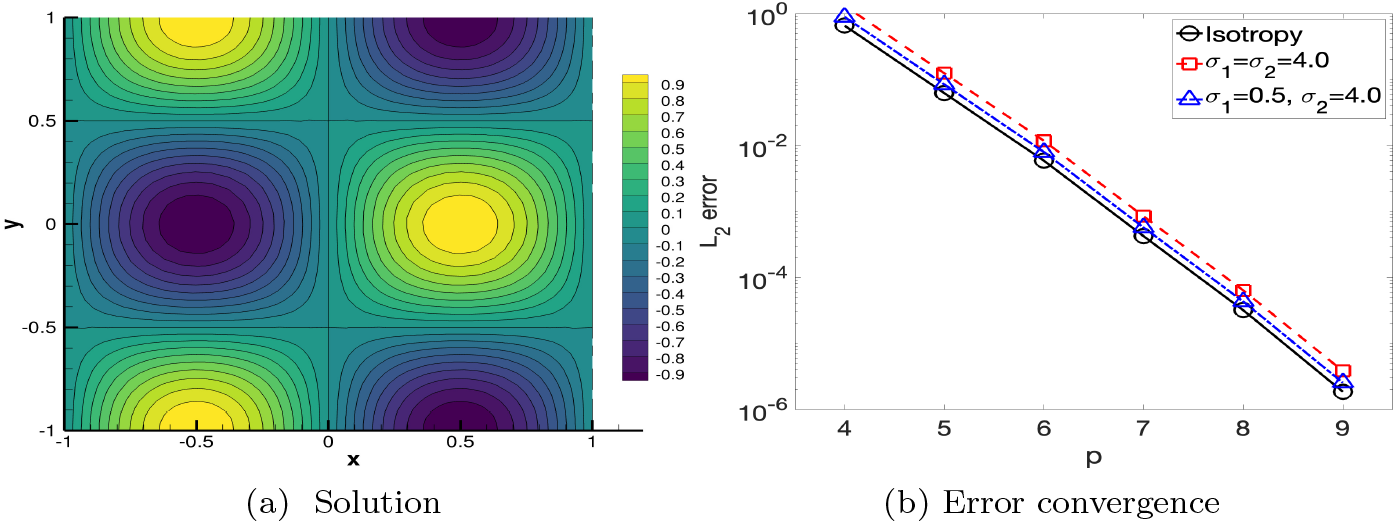
Anisotropic Helmholtz solver for a test problem with (a) the solution, and (b) error convergence for various *σ*_1_ and *σ*_2_.

## 3 Results

The computational results were obtained by numerically solving the proposed multidimensional neural spike propagation model in Eqs. (54) to (55). The numerical scheme is given as a weak formulation with moving frames in the context of the continuous Galerkin formulation, as presented in Eqs. 56 with the boundary conditions in Eqs. (60) to (61). The scheme was implemented in the open-source spectral/hp element framework, called *Nektar++* [36]. 1D data are extracted along a line from 2D simulations to analyze the propagation along the fiber, particularly regarding arriving time and conduction velocity. Moreover, this work introduced the *arrival time map* (i.e., *time map*) denoted by 𝒯(**x**) for **x** ∈ Ω, for the comparison and analysis of conduction speed and synchronization. The time map, corresponding roughly to the time when the neural spike is excited, is computed as follows [17]:

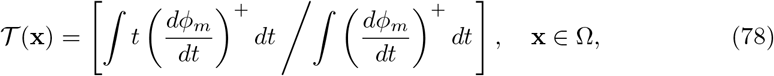

where the superscript + denotes the maxmod function defined by

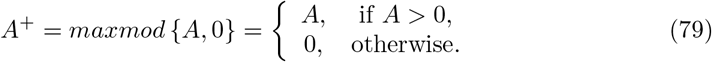

In this paper, the width of the intracellular domain, regarded as a fiber bundle with identical fibers as a small fascicle, is primarily 0.01 cm in most cases. Considering that the fiber radius is approximately 2 *µ* m, the corresponding bundle with a width of 0.01 cm contains around a thousand neural fibers for a packing ratio of 0.5. In the proposed mathematical model, ephaptic coupling between fiber bundles is only effective when ephaptic coupling within each fiber bundle is ignored. Thus, the conduction velocity of the neural spike along the fiber bundle is approximately the same as that of a single fiber bundle. This conduction velocity contrasts the fact that the conduction velocity in an ephaptically coupled fiber bundle slows [43, 49]. The mathematical models in Eqs. (54) to (55) can be further modified to include this slowing effect in a fiber bundle. However, this model was not considered in this paper because it is beyond the scope of work.

### A. Single fiber bundle

This work considers a single fiber bundle embedded in a 2D plane, as presented in Fig. 7a. The domain of the fiber bundle is given as a rectangular region (*x, y*) ∈ [*x*_0_, *x*_0_ + *w*] × [*y*_0_, *y*_0_ + *L*], where *w* denotes the fiber width at 0.01 cm, and *L* represents the fiber length varying from 1 to 4 cm. Each node and myelin length is the same at 0.01 and 0.2 cm, respectively. Temperature is 24°*C*, and conductivity ratio *rho* is 1. The neural spike is initiated at the left-most node for the duration of 0.01 ms with a continuous external 0.4 mV stimulus. Without myelin, the shape of the neural spike is almost linear in the depolarization and repolarization phases, as depicted in Fig. 7c, coincident with the findings of previous studies [18, 43]. The conduction velocity in an unmyelinated axon is close to 2.5 m/s, consistent with the previously known value [58].

**Figure 7:**
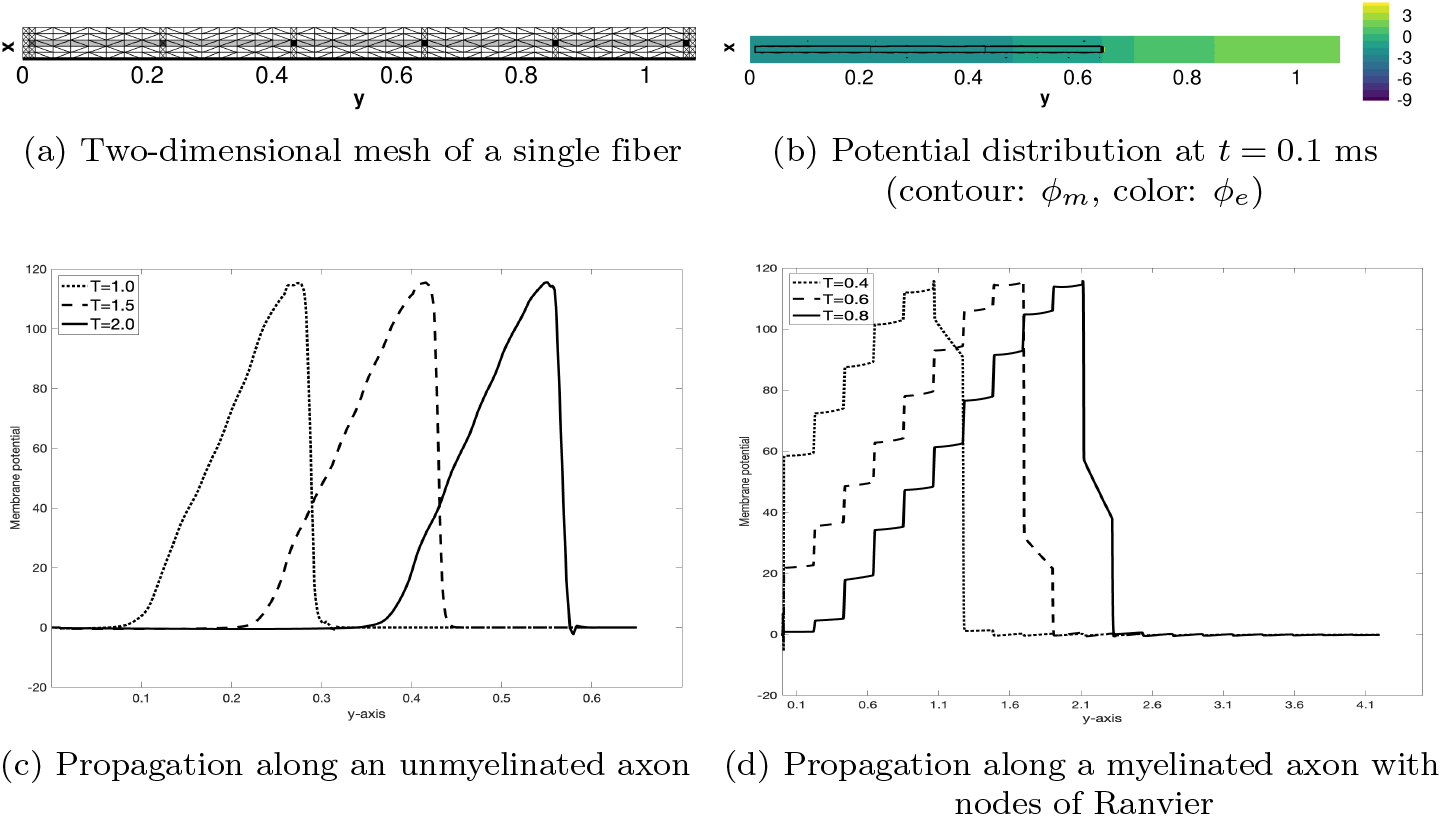
Single-fiber propagation and comparison of neural spikes in the neural fiber in unmyelinated and myelinated axons with nodes of Ranvier. The unit of time is milliseconds and the length is centimeter. (a) The myelin sheath is marked gray, and the four nodes of Ranvier are slightly darker, (*x, y*) = [0.02, 0.03] ×[0.02+*k* ∗0.2, 0.03+*k* ∗0.2] for 0 ≤ *k* ≤ 3. *x*_0_ = 0.01, *y*_0_ = 0.01, *x*_*L*_ = 0.0, *x*_*R*_ = 0.05, *y*_*L*_ = 0.0, *y*_*R*_ = 0.65, (b) Potential distribution at *t* = 0.4 ms initiated from the leftmost node. contour: *ϕ*_*m*_, color: *ϕ*_*e*_. (c) Simulated on a fiber of length 0.65 cm. (d) Simulated on the fiber with 20 nodes and a length of 4.1 cm, where *T* = 24°*C, ρ* = 1.0, and *r*_*b*_ = 0.1 cm.

The myelinated sheath increases the neural spike duration by more than 10 times larger than that of the unmyelinated axon. The wavelengths in the unmyelinated and myelinated axons with the nodes of Ranvier are approximately 0.2 cm and 2.2 cm, respectively. Moreover, the conduction velocity in the myelinated axon with nodes of Ranvier is also increases by more than 10 times that of the unmyelinated axon (i.e., approximately 25 m/s) in accordance with the measured quantity of a frog motor axon [53] and the corresponding computational simulations of previous studies [18, 43]. In Fig. 7d, the most striking feature of the neural spike for the myelinated axon is the dramatic potential change at the nodes of Ranvier, in contrast to slower changes in the myelin sheath. The reaction of the ion channels causes this staircase repolarization phase only at the nodes of Ranvier. The highest magnitude of the action potential for unmyelinated and myelinated axons remains approximately the same.

Furthermore, Fig. 8 displays the time sequential distribution of the extracellular potential (*ϕ*_*e*_) distribution perpendicular to the propagation direction, i.e., along the *x*-axis. The membrane potential increases for the initial depolarization (*t* = 0.05 ms). Accordingly, the extracellular potential inside the axon increases (*t* = 0.075 ms). After the extracellular potential reaches a certain threshold, for example, approximately around 10 mV for a fiber radius of 0.1 cm (*t* = 0.1 ms), the extracellular potential becomes approximately zero when the membrane potential is approximately 60 mV (*t* = 0.12 ms). The extracellular potential becomes negative when the membrane potential is between 80 and 120 mV (*t* = 0.15 ms). Finally, it returns to zero when the membrane potential reaches the maximum (*t* = 0.18 *ms*). This extracellular potential fluctuation is consistent with the mathematical predictions for the oscillation between hyperpolarization and depolarization deduced by Markin in the 1970s [29–32].

**Figure 8:**
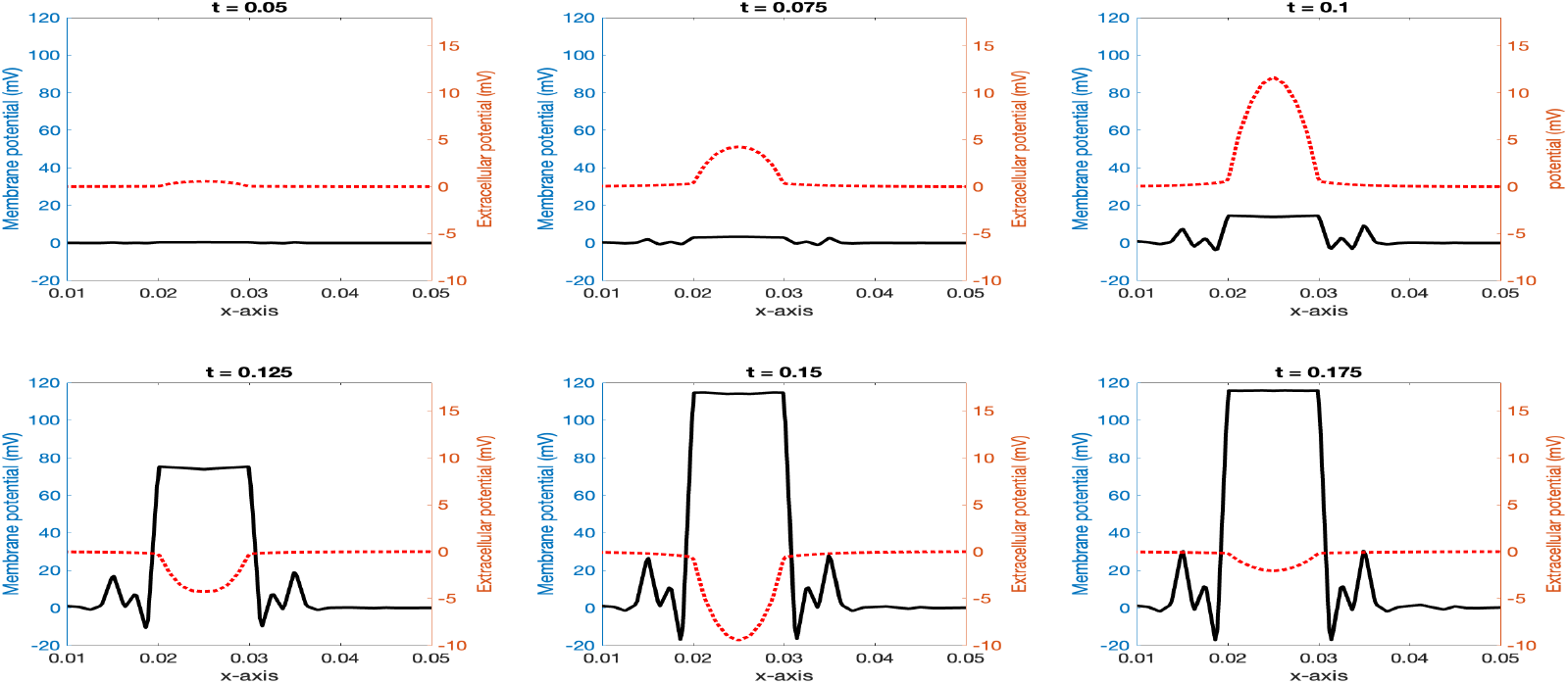
Distribution of the membrane potential (*ϕ*_*m*_) and extracellular potential (*ϕ*_*e*_) perpendicular to the propagation direction (along the *x*-axis). Time *t* is in milliseconds. Measured at a node of Ranvier, wherew *T* = 24°*C, ρ* = 1.0, and *r*_*b*_ = 0.1 cm.

Moreover, Fig. 9 displays the conduction velocity changes according to the conductivity ratio *ρ* variation between the intracellular and extracellular space. In addition, Fig. 9a displays the variations of the instantaneous conduction velocity of a single neural spike initiated at the left-most node using the instantaneous velocity from the arrival time [17]. The contribution of the extracellular potential is proportional to the inverse of the square of the distance from the nodes of Ranvier, as deduced from Eq. (35), or deduced in [48, 49]. Thus, the instantaneous velocity varies from 10 ∼ 60 (m/s) for the corresponding value of *ρ* as {1, 2, 4, 8, 12}.

**Figure 9:**
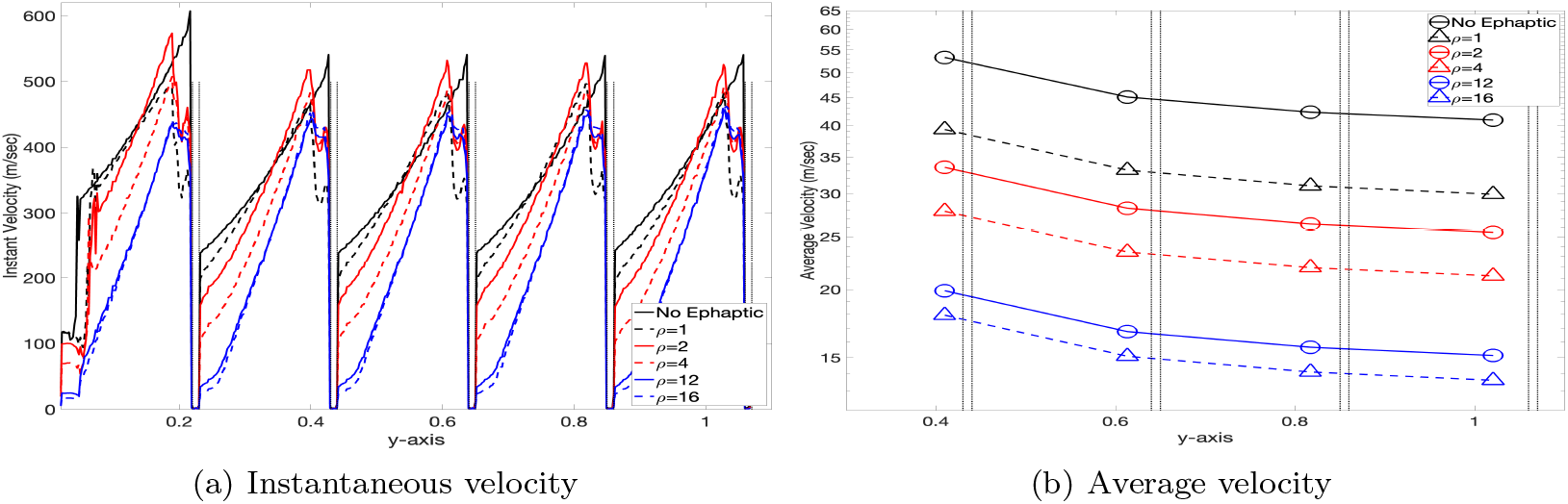
Conduction velocity along a single fiber bundle. The vertical dotted black line corresponds to the boundaries of the nodes of Ranvier, where *ρ* denotes the conductivity ratio between the intracellular and extracellular domains, *T* = 24°*C*, and *r*_*b*_ = 0.1 cm.

The average velocity of the neural spike propagation along the myelinated axon with the nodes of Ranvier is obtained by averaging the instantaneous velocity, as illustrated in Fig. 9b. The average velocity depends on *ρ*, coincident with the result from [43].

Moreover, the average conduction velocity is approximately proportional to the square root of *ρ*. For example, the average velocity with *ρ* = 1 is between 36 and 52 m/s, approximately three times larger than the average velocity for *ρ* = 12 when the conduction velocity is between 13 and 20 m/s.

Moreover, Table 3 demonstrates that the average conduction velocity depends on the fiber radius and temperature. In the mathematical model of Eq. (28) using the approximation of Eq. (35), the fiber radius does not significantly change the conduction velocity for a single fiber bundle. The induced extracellular potential primarily causes a slight change in the conduction velocity. The proposed mathematical model does not consider the direct conduction velocity change in *I*_*ion*_ by the fiber radius, as elaborated after Eq. (35), causing this non-dependency. In contrast, the increase in temperature directly yields a dramatic increase in the conduction velocity. Temperature is a critical component in the function *Ƶ* in Eq. (12), affecting the threshold of ion channels. This computational simulation is consistent with the physiological observation in [42].

**Table 3:** Conduction velocity dependency along a single fiber. For, the conduction velocity versus the fiber radius, *T* = 24°*C*. For the conduction velocity versus temperature, *r*_*b*_=0.1 cm. *ρ*=1.0.

|  |  |  |  |  |  |
| --- | --- | --- | --- | --- | --- |
| Fiber radius (cm) | 0.01 | 0.1 | 0.2 | 0.4 | 0.8 |
| Conduction velocity (m/s) | 29.0 | 29.3 | 29.5 | 29.5 | 29.4 |
| Temperature ($^\circ C$) | 24 | 28 | 32 | 36 | 40 |
| Conduction velocity (m/s) | 29.3 | 45.3 | 70.3 | 108.8 | 168.3 |

Table 3 reveals that the conduction velocity linearly increased up to 30°*C* but increased more rapidly for higher temperatures when *ρ* = 1 [43].

### B. Fiber bundles with aligned nodes of Ranvier

This work considers two straight and parallel fiber bundles with aligned node of Ranvier, as depicted in Fig. 10a. The two fiber bundles with the same width (0.01 cm) are aligned along the *y*-axis with a gap of *w* = 0.01 cm between them along the *x*-axis. The neural spike propagation is initiated at the nodes at *y* ∈ [0.01, 0.02] for the first fiber bundle in *x* ∈ [0.01, 0.02] and the second in *x* ∈ [0.03, 0.04].

**Figure 10:**
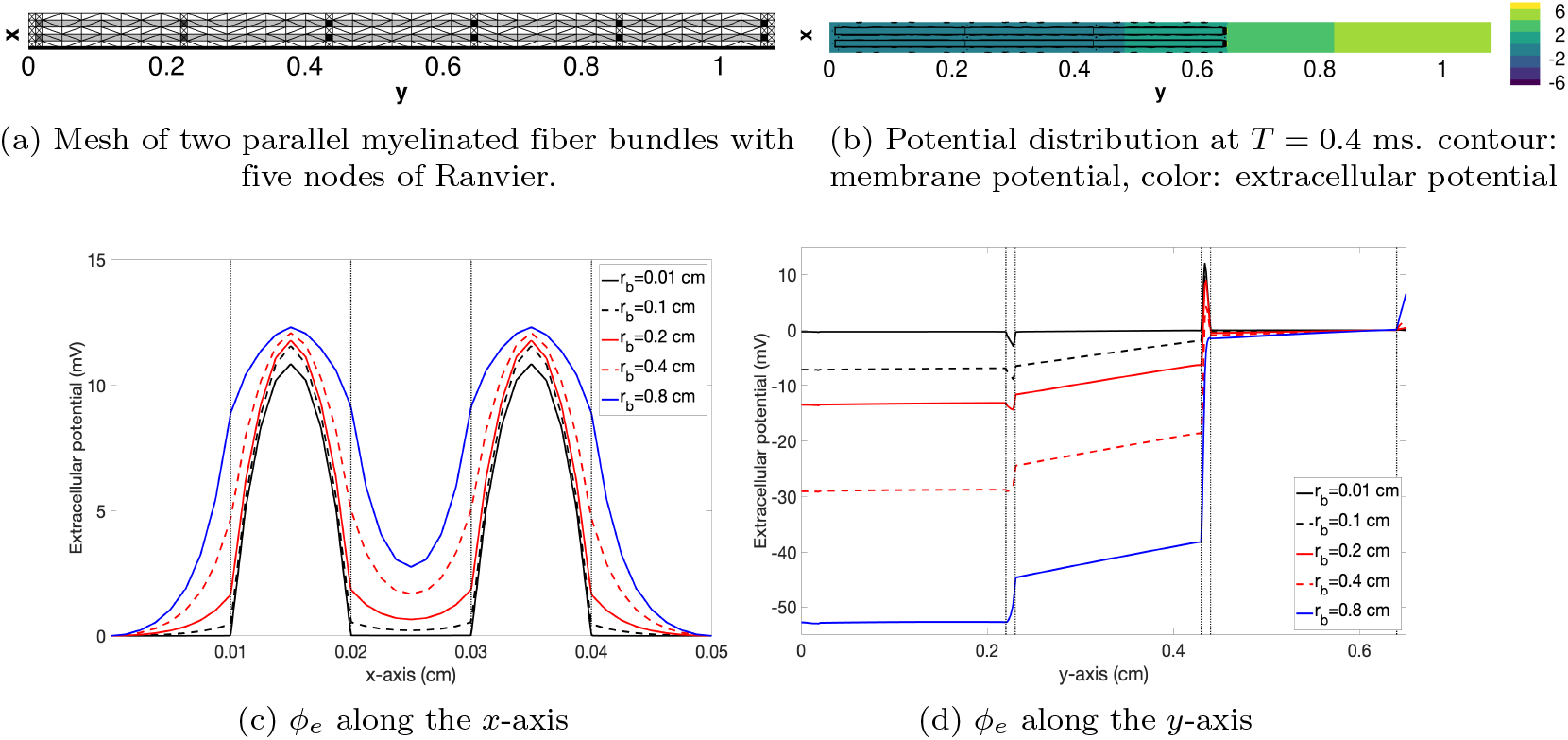
Extracellular potential distribution along each axis for two fiber bundles with aligned nodes of Ranvier. The first fiber bundle is in *x* ∈ [0.01, 0.02] and the second is in *x* ∈ [0.03, 0.04], where *T* = 24°*C* and *ρ* = 1.0.

First, the two fiber bundles are initiated simultaneously in the first node of each fiber bundle. The propagation along the first and second fiber bundles is approximately the same, with an arrival time error of 10^−5^ ms. Similar to the propagation in a single fiber bundle, the strength of the extracellular potential *ϕ*_*e*_ also increases as the fiber bundle radius increases, as illustrated in Fig. 10c. For example, the extracellular potential between the fiber bundles with a radius 0.8 cm is approximately 3 mV when the extracellular potential in each fiber bundle is approximately 9 mV. This potential is significantly greater than the extracellular potential between the fiber bundles with a radius of 0.1 cm, approximately 0.216 mV. Moreover, Fig. 10d demonstrates that the extracellular potential changes drastically after depolarization where the magnitude is approximately proportional to the fiber bundle radius. The increase in the propagation velocity caused by the fiber bundle radius is relatively insignificant to other factors, such as *ρ* and temperature.

In Fig. 11, the propagation in the myelin sheath of the two fiber bundles is faster than that in the single fiber bundle, regardless of the conductivity ratio (*ρ*) and fiber bundle radius (*r*_*b*_). The difference in the conduction velocity between these two fiber bundles is small for *r*_*b*_ = 0.01 cm, as illustrated in Fig. 11b; thus, this is likely caused by ephaptic coupling effects. Contrary to the conduction velocity in the myelin sheath, the conduction velocity in the nodes of Ranvier depends on the ephaptic coupling effects and the conductivity ratio *ρ*. Table 4 demonstrates that the conduction velocity at the nodes of the two fiber bundles is slower than that at the nodes of the single fiber bundle for a smaller *ρ*, less than about *ρ* = 3.0. The conduction velocity at the nodes of Ranvier of two fiber bundles with a *ρ* value larger than 3.0 is faster than that for a single fiber bundle. Hence, the collective average velocity in two fiber bundles is slower than that of a single fiber bundle for a *ρ* value less than 3.0. If *ρ* is higher than 3.0 with stronger ephaptic coupling effects, the propagation in two fiber bundles is faster than that of a single fiber bundle. The slowing of the propagation in the fiber bundles by ephaptic coupling for a low *ρ* value agrees with previous 1D studies [19, 43, 49]. However, the increase in speed of the conduction velocity for two fiber bundles for a large value of *ρ* was unknown.

**Table 4:** Conduction velocity at the fourth node and the total average velocity in the myelin and nodes for the propagation 1) without ephaptic coupling (no ephaptic), 2) with two fiber bundles, and 3) with a single fiber bundle. (*) Difference is measured as the conduction velocity of two fibers minus the conduction velocity of a single fiber bundle.

| $\rho$ (conductivity ratio) | | 1 | 2 | 4 | 8 | 12 |
| --- | --- | --- | --- | --- | --- | --- |
| Node velocity (m/s) | No ephaptic | 5.366 |  |  |  |  |
|  | Two fiber bundles | 3.661 | 2.804 | 2.096 | 1.543 | 1.572 |
|  | Single fiber bundle | 3.723 | 2.795 | 2.068 | 1.542 | 1.481 |
|  | Difference (*) | -0.0618 | 0.0089 | 0.028 | 0.0007 | 0.0918 |
| Avg. velocity (m/s) | No ephaptic | 36.188 |  |  |  |  |
|  | Two fiber bundles | 26.005 | 22.246 | 18.637 | 15.341 | 13.482 |
|  | Single fiber bundle | 26.245 | 22.352 | 18.614 | 15.213 | 13.272 |
|  | Difference (*) | -0.24 | -0.11 | 0.023 | 0.13 | 0.21 |

**Figure 11:**
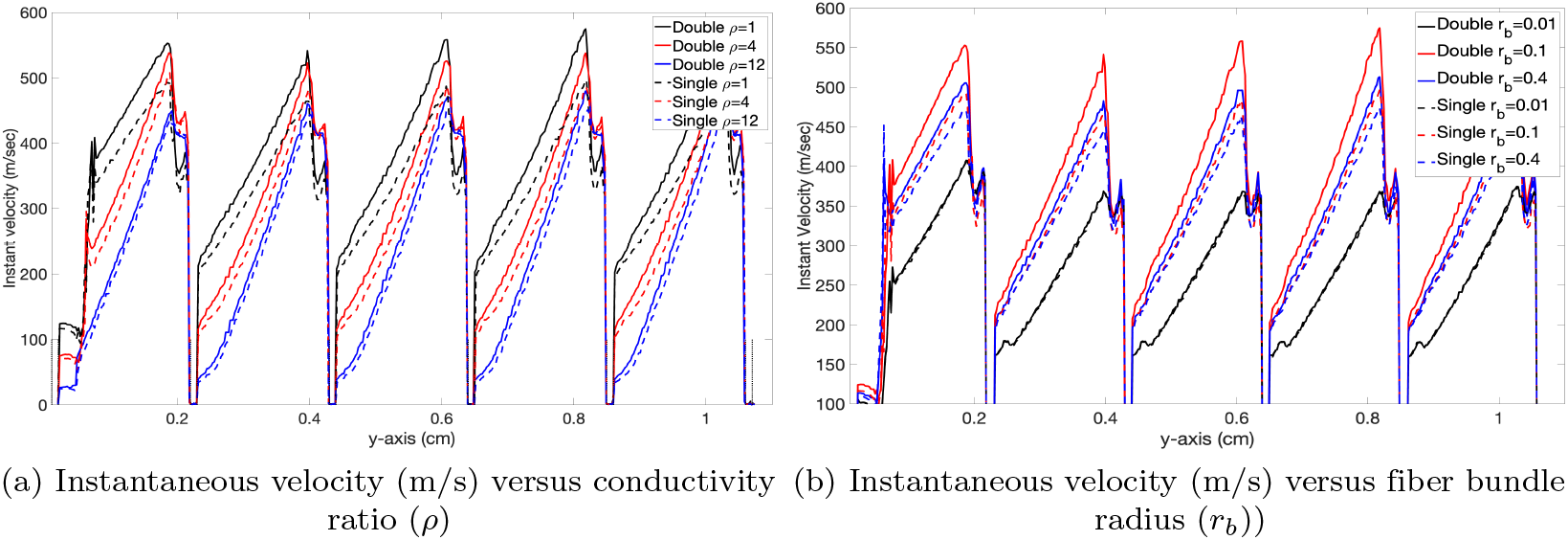
Instant conduction velocity in two fiber bundles with simultaneous initiations. (a) Instant velocity along the first fiber bundle for various conductivity ratio *ρ. r*_*b*_ = 0.1 cm. *Double* and *single* correspond to the velocity for two and a single fiber, respectively. (b) Instant velocity along the first fiber bundle for various fiber bundle radius *r*_*b*_ (*ρ* = 1.0). The two fiber bundle’ radii are 0.01 cm, and the gap between them is 0.01 cm (*T* = 24°*C*).

This work considers the propagation of the two fiber bundles with a delayed initiation in one fiber bundle, where the initiation of the second fiber bundle is delayed without loss of generality. Both Fig. 12 and Table 5 demonstrate the effects of ephaptic coupling on unsynchronized propagation, agreeing with previous mathematical and computational studies [43, 48]. in Fig. 12a, synchronization continuously occurs between the two fiber bundles when the initiation delay is less than about 0.1 ms. This time is approximately when the neural spike in the first fiber leaves the first node (*y* ∈ [2.2, 2.3]) and moves into the second myelin (*y* ∈ [2.3, 4.3]).

**Table 5:** Synchronization velocity and relative synchronization velocity defined in Eq. (80) for the two fiber bundles with aligned nodes of Ranvier. The initiation delay occurs in the second fiber bundle of *y* =∈ [0.03, 0.04]. The two fiber bundles have a width of 0.01 cm and length of 2.2 cm with a gap of 0.01 cm between them, where *ρ* = 1, *r*_*b*_ = 0.1 *cm*, and *T* = 24°*C*.

| Delay (ms) | 0.01 | 0.02 | 0.04 | 0.08 | 0.1 | 0.11 |
| --- | --- | --- | --- | --- | --- | --- |
| Sync. velocity (s/cm) | 2.48 | 4.76 | 7.55 | 4.77 | 0.86 | -0.93 |
| Relative sync. velocity (%/cm) | 25.8 | 24.5 | 19.8 | 6.3 | 0.9 | -0.9 |

**Figure 12:**
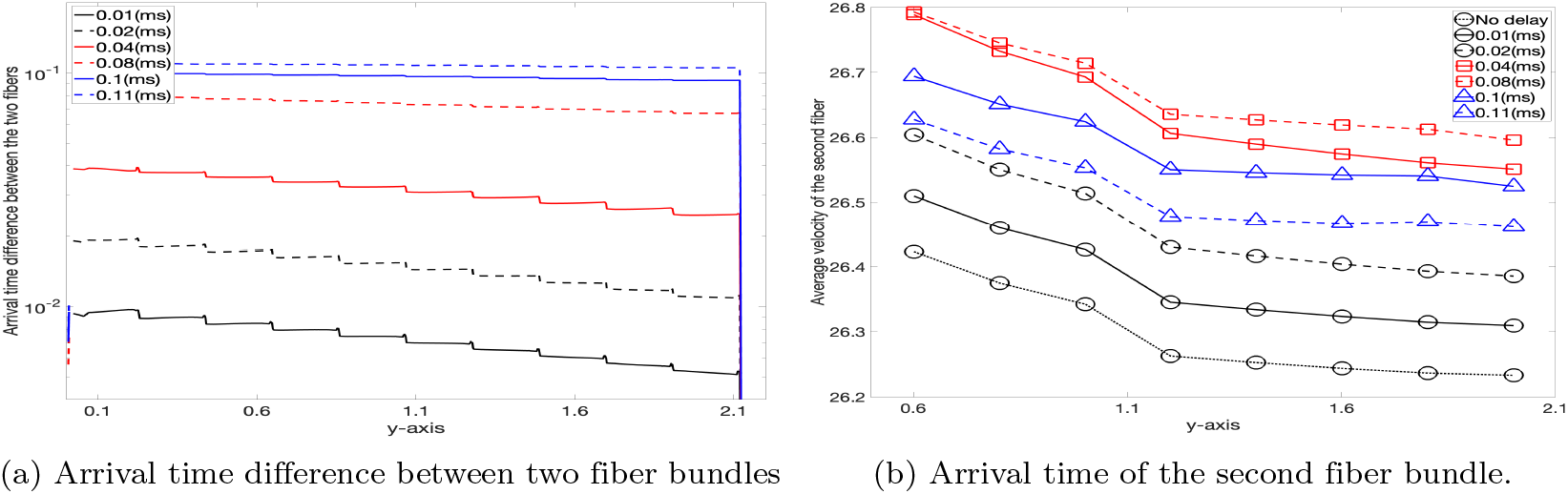
Propagation along two fiber bundles with an initiation delay. (a) Arrival time difference between the two fiber bundles when the second fiber bundle is initiated with various delays. (b) Conduction velocity of the second fiber bundle increases for synchronization, whereas the conduction velocity of the first fiber bundle remains about the same.

Moreover, Fig. 12b confirms that synchronization is caused by the increased velocity of the delayed neural spike in the second fiber bundle while the velocity of the first fiber bundle remains approximately the same. The average velocity of the second fiber is at maximum when the initial delay is 0.04 ∼0.08 ms and is at minimum for the initial delay of more than 0.1 ms or less than 0.01 ms. The synchronization velocity and ratio are defined as follows to quantify the rate of each synchronization:

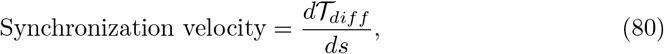

where 𝒯_*diff*_ denotes the arrival time difference between the fiber bundles, and *s* represents the parameter of the propagation distance. Similarly, the relative synchronization velocity is defined by dividing Eq. (80) by the maximum arrival time difference. Table 5 reveals that the synchronization velocity is linearly proportional to the magnitude of the initial delay, particularly up to a delay of 0.05 ms. This synchronization proportionality to the initial delay provides a constant synchronization ratio when the initial delay is less than 0.05 ms.

This work also considers three fiber bundles with aligned nodes of Ranvier, as illustrated in Fig. 13. The configuration of the fiber bundles is the same as the case for two fiber bundles. The three fiber bundles are aligned along the *y*-axis, and each domain is defined as follows: *y* ∈ [0.01, 0.02], *y* ∈ [0.03, 0.04], and *y* ∈ [0.05, 0.06 in that order. The neural spike initiate at the same location in the three fibers but with a slight delay in the second fiber bundle. The extracellular potential distribution for the initial delay of 0.04 ms in the second fiber bundle is displayed in Fig. 13c. The synchronization in the three fiber bundles is slightly faster than for the two for a shorter delay of approximately less than 0.05 ms, as presented in Table 6. However, synchronization is significantly slower or fails when the delay is longer than 0.08 ms, which is less than that for the two fiber bundles. In this paper, synchronization in similar conditions is reported for a delay of 0.5 ms in [43] for a packing ratio of 0.25, in contrast to a ratio of 0.5 for the configuration of the three fiber bundles. The packing ratio represents the ratio between the extracellular and intracellular space in a fiber bundle. This mechanism is likely due to the symmetric extracellular potential distribution. This potential distribution increases the synchronization within a shorter delay but lowers the delay threshold for synchronizing three fiber bundles.

**Table 6:** Absolute and relative synchronization velocity defined in Eq. (80) for the three fiber bundles with aligned nodes of Ranvier. The initiation delay occurs in the second (middle) fiber bundle of *y* =∈ [0.03, 0.04]. The threes fiber bundles have a width of 0.01 cm and length of 1.1 cm with a gap of 0.01 cm between them, where *ρ* = 1, *r*_*b*_ = 0.1 cm, and *T* = 24°*C*.

| Delay (ms) | 0.01 | 0.02 | 0.04 | 0.08 | 0.1 | 0.12 |
| --- | --- | --- | --- | --- | --- | --- |
| Sync. velocity (s/cm) | 2.45 | 5.32 | 8.82 | 2.93 | -0.23 | -4.83 |
| Relative sync. velocity (%/cm) | 26.3 | 28.0 | 22.8 | 3.8 | -0.2 | -4.1 |

**Figure 13:**
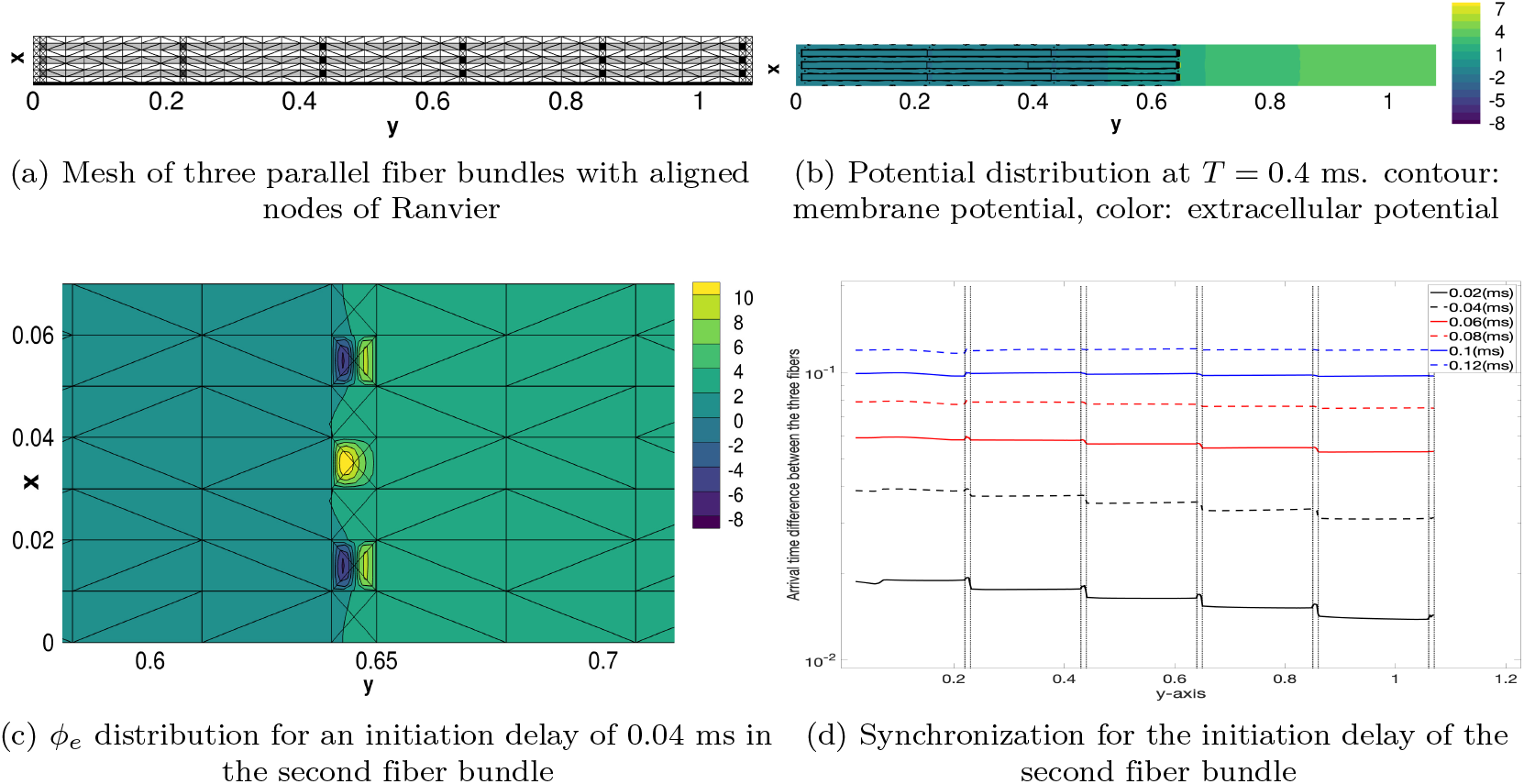
Propagation in the three neighboring fiber bundles. The first, middle, and third fiber bundles are located in *x* ∈ [0.01, 0.02], *x* ∈ [0.03, 0.04] and *x* ∈ [0.05, 0.06], respectively. (a) Mesh of the three fiber bundles with the same thickness of 0.01 cm and a gap of 0.01 cm between them. (b) Distribution of the extracellular space for a delayed initiation of the middle fiber bundle. (c) Synchronization of neural spike propagation in the three-parallel fiber bundles for an initiation delay in the second fiber bundle.

### C. Interference: Opposite-traveling neural spikes

The first simulation, which is only available in a multidimensional model, is the interference between the traveling spikes in different directions. As an example of interference, the neural spike propagations in two parallel but opposite-propagating fiber bundles are simulated; one propagation in the normal (orthodromic) direction along one fiber bundle and the other in the opposite (antidromic) direction along the other fiber bundle. The configuration of the two fiber bundles is the same as that in the previous section but with different initiation nodes. The first and second fiber bundles initiate from the left-most (*y* ∈ [0.01, 0.02]) and right-most (*y* ∈ [1.08, 1.09]) nodes, respectively. Further, Figs. 14b to 14d display the 1D distribution of the membrane and extracellular potential at various time before and after the two spikes are closest.

**Figure 14:**
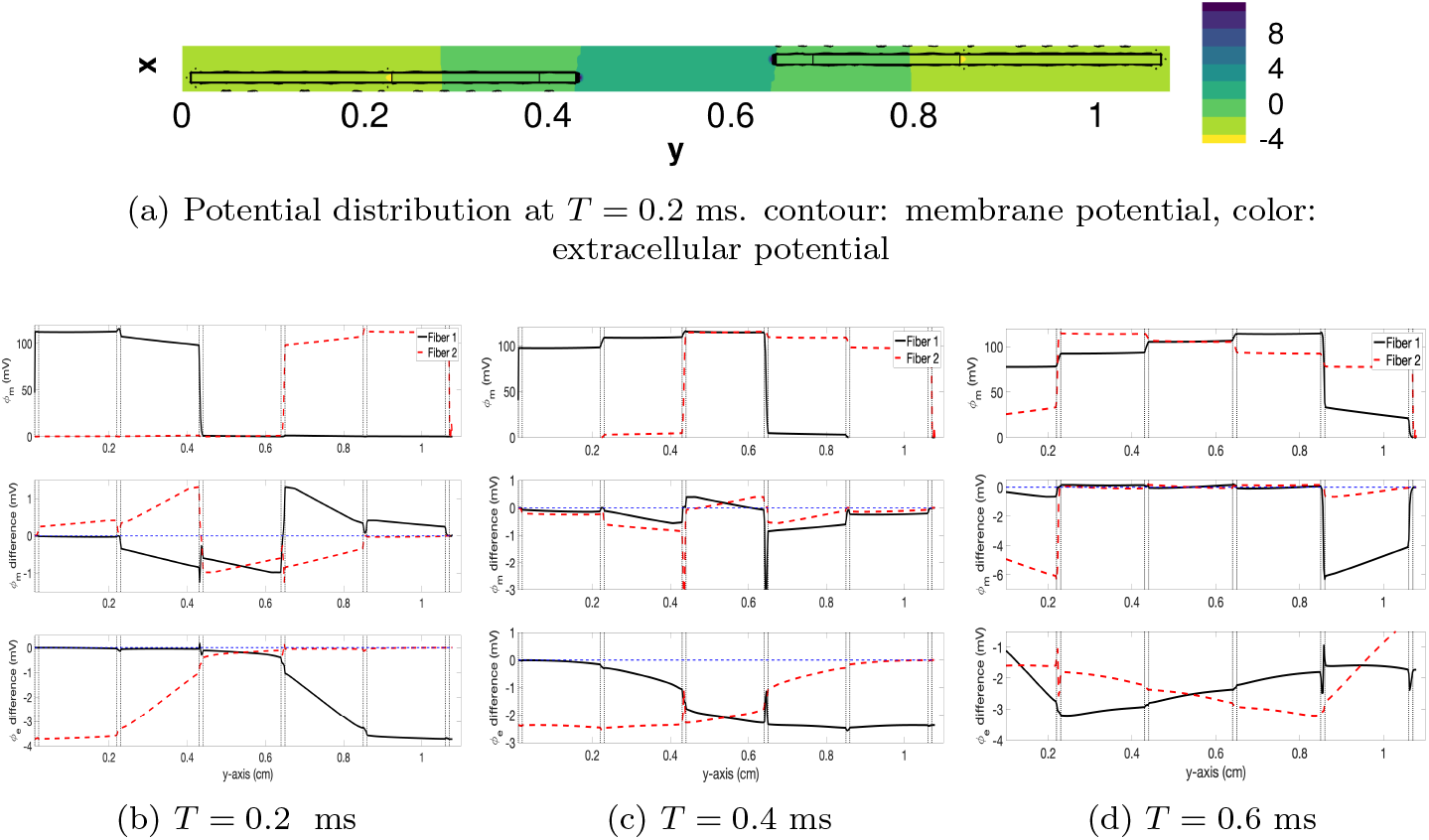
Propagation along the two fiber bundles with six nodes of Ranvier in the normal and opposite directions. The mesh is the same as in Fig. 10a. The first fiber in *x* ∈ [0.01, 0.02] is initiated at *y* ∈ [0.01, 0.02] and the second fiber in *x* ∈ [0.03, 0.04] is simultaneously initiated at *y* ∈ [1.06, 1.07], where *ρ* = 1, *r*_*b*_ = 0.1 cm, and *T* = 24°*C*.

The membrane potential in one fiber bundle does not directly affect the other because each fiber domain is separated. However, the extracellular potential distribution in the extracellular space is abruptly changed by interference. The interference has two distinctive patterns, depending on when the two neural spikes are closest; the first is when the two spikes are closest at a node and the second is when the two spikes are closest at the myelin sheath.

The first case can be represented by two opposite-traveling fibers with an even number of nodes, as depicted in Fig. 10a. The two neural spikes are closest in the middle myelin when the spikes are initiated from opposite ends. Moreover, Fig. 14 displays the membrane and extracellular potential distribution of two opposite-traveling neural spikes. Without opposite-traveling spikes, the extracellular potential of the first fiber bundle is zero ahead of the depolarizing area. If the opposite-traveling spike occurs ahead of the depolarizing area, the extracellular potential distribution follows the same distribution as that of the second fiber bundle (Fig. 14b). This deformation of the extracellular potential ahead of the depolarization area generates an additional electric current to modify the electric current in the fiber. The extracellular potential gradients are presented in *y* ∈ [0.6, 0.8], [0.2, 0.6], and [0.0, 0.2] for T=0.2, 0.4, and 0.6 ms, respectively. The extracellular potential gradients caused by the interference seem to propagate backward. This extracellular potential gradient induces the corresponding membrane potential changes, as depicted in Figs. 14b to 14d.

The second case is represented well by two opposite-traveling fibers with an odd number of nodes. When the spikes are initiated from opposite ends, the two neural spikes are closest to a node of Ranvier, as illustrated in Fig. 15b. Similar to fibers with an odd number of nodes, the extracellular potential gradient by interference emerges in *y* ∈ [0.6, 0.8] and [0.4, 0.8], and approximately 0.3 for *T* = 0.3, 0.5, and 0.7 ms, respectively. Compared to the fiber with an even number of the nodes of Ranvier, the extracellular potential deformation is more substantial; thus, the membrane potential change in each fiber is more significant. Nevertheless, the deformation region seems smaller than that for fibers with an even number of nodes of Ranvier. The effect of this kind of interference in the fiber with an odd number of nodes of Ranvier appears reasonable because the two neural spikes remain closest at the neighboring nodes of Ranvier, with a stronger but narrower interference with the extracellular potential of other spikes.

**Figure 15:**
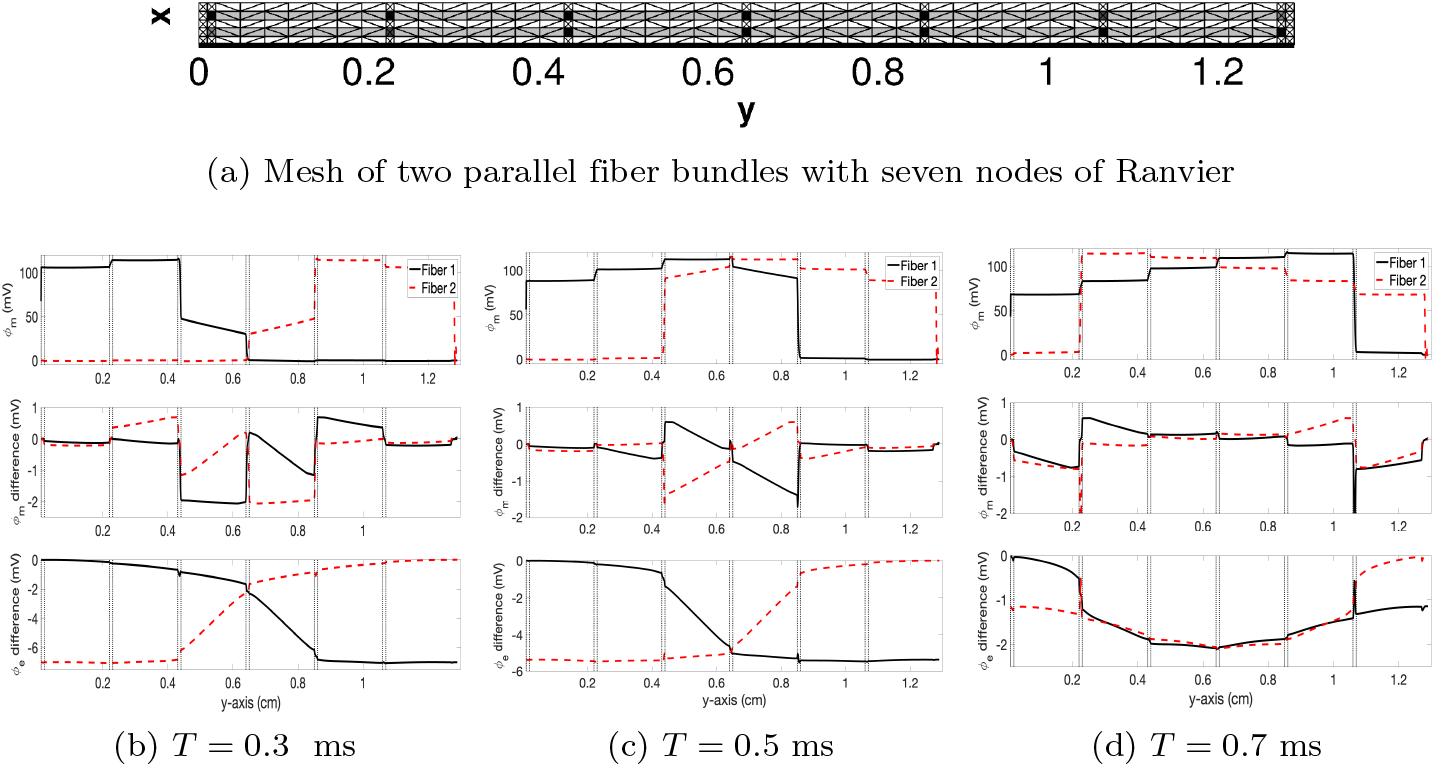
Propagation along the two fiber bundles with seven nodes of Ranvier in the normal and opposite directions. The mesh is the same as in Fig. 10a. The first fiber bundle in *x* ∈ [0.01, 0.02] is initiated at *y* ∈ [0.01, 0.02], and the second fiber bundle in *x* ∈ [0.03, 0.04] is simultaneously initiated at *y* ∈ [1.27, 1.28], where *ρ* = 1, *r*_*b*_ = 0.1 cm, and *T* = 24°*C*.

Ephaptic coupling of traveling neural propagations depends on when the two neural spikes are closest. Thus, the configuration of the fiber bundle or the initiation delay can affect the interference magnitude. For example, a variation in the delay of the second fiber bundle can generate changes in the propagation velocity of the first fiber by interference, significantly modifying its arrival time. In addition, Fig. 16a confirms that the conduction velocity of the first fiber bundle is approximately proportional to the delay of the second fiber, up to 0.05 ms. Moreover, the fiber radius also affects the interference because the change in the fiber bundle radius yields a change in the magnitude of ephaptic coupling. Further, Fig. 16b confirms that the larger ephaptic coupling effect does not always yield a more significant change in the arrival time and conduction velocity. Significant slowing and greater interference occur at *r*_*b*_ = 0.2 cm.

**Figure 16:**
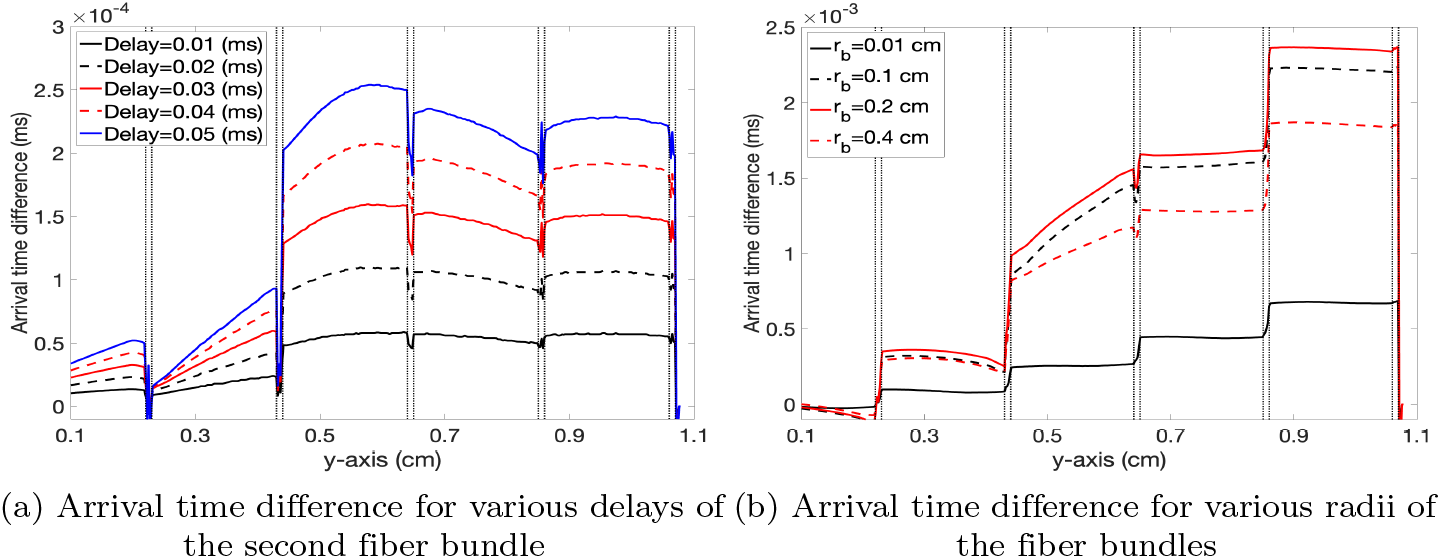
Arrival time difference of the first fiber bundle against (a) the delay time for the second fiber bundle and (b) the fiber bundle radius. The fiber bundles have an even number of nodes. The arrival time difference is computed as the arrival time for two opposite-traveling fiber minus the arrival time for a single fiber. The mesh is the same as in Fig. 10a. The first fiber bundle in *x* ∈ [0.01, 0.02] is initiated at *y* ∈ [0.01, 0.02], and the second fiber bundle in *x* ∈ [0.03, 0.04] is initiated at *y* ∈ [1.06, 1.07], where *ρ* = 1, *r*_*b*_ = 0.1 cm, and *T* = 24°*C*.

### D. Asynchronizing configuration: Misaligned nodes of Ranvier

The second available case using the multidimensional bidomain model is neural spike propagation in the fiber bundles with misaligned nodes of Ranvier. This work considers two parallel fibers with misaligned nodes of Ranvier with the first and second fiber bundle in *x* ∈ [0.01, 0.02] and *x* ∈ [0.03, 0.04], respectively, as illustrated in Fig. 17a and 17b. The nodes of Ranvier of the first fiber bundle are not aligned with those of the second fiber bundle but alternate every 0.1 cm, except the first and last nodes. For example, the third nodes of the first and second fiber bundles appear at *y* ∈ [0.23, 0.24] and [0.33, 0.34], respectively. The total length of the myelin and fiber is the same for the two fibers.

**Figure 17:**
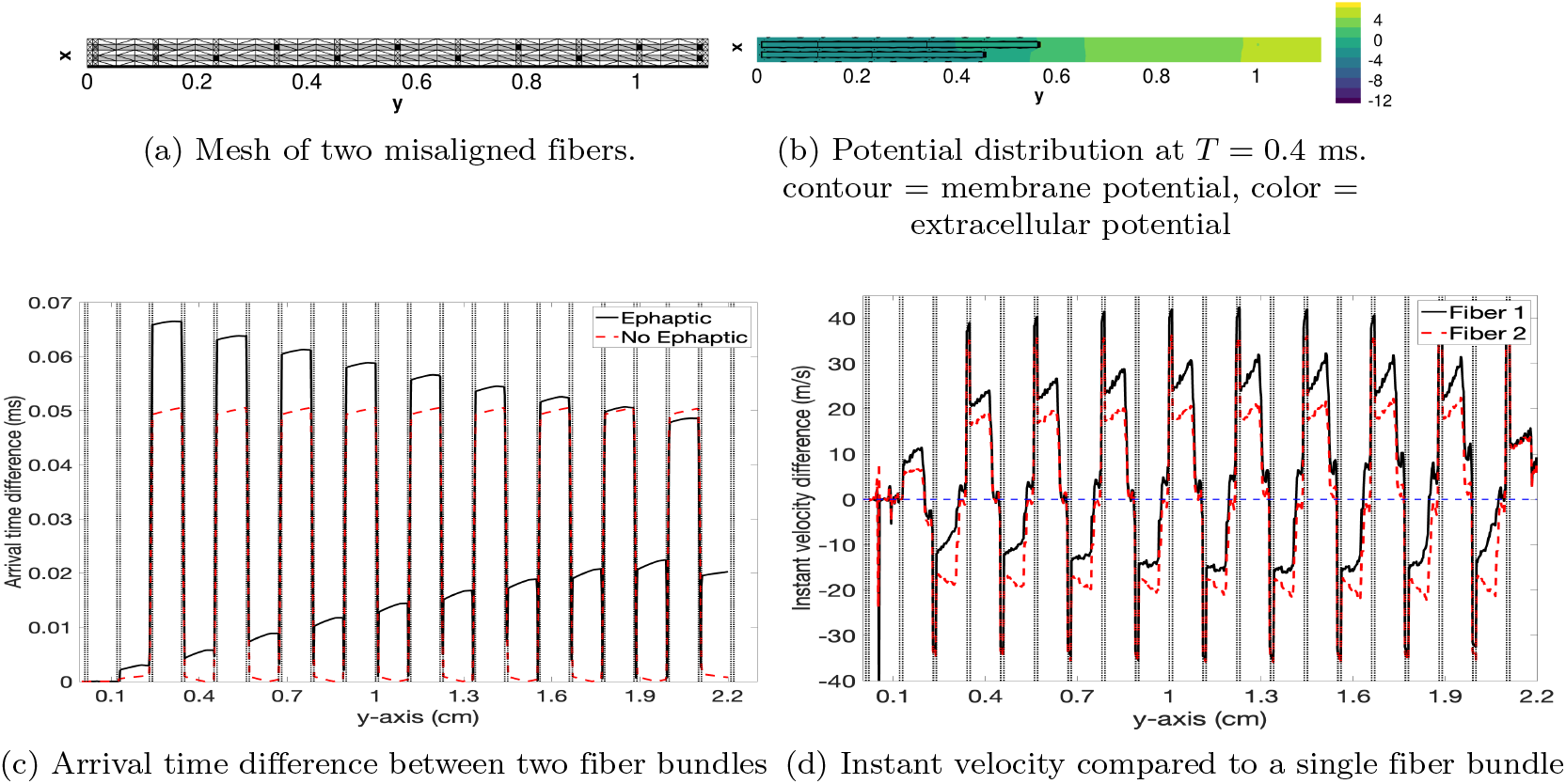
Arrival time and instant velocity differences in fiber bundles with misaligned nodes of Ranvier. The fiber bundle configuration is the same as before, except for the misaligned and alternating nodes of Ranvier. (a) Mesh of the misaligned nodes of Ranvier (dark square). Fiber bundles 1 and 2 belong to the domain of *x* ∈ [0.010.02] and *x* ∈ [0.03, 0.04], respectively. (b) Membrane and extracellular potential distribution at *T* = 0.4 ms. (c) Significant arrival time difference (0.0202 ms) via ephaptic coupling at the final node of Ranvier, compared to the negligible time difference (7.63 × 10^−4^ ms) at the final node of Ranvier via nonephaptic propagation. (d) Instant velocity difference for each fiber bundle displays complex velocity changes via ephaptic coupling in fiber bundles with misaligned nodes, where *ρ* = 1, *r*_*b*_ = 0.1 cm, and *T* = 24°*C*.

If two fiber bundles are initiated simultaneously at the first nodes in each fiber bundle, without the ephaptic coupling effect, the neural spike reaches its final node simultaneously, or with a small delay as small as 7.63 × 10^−4^ ms for a fiber with a length of 2.22 cm with 10 misaligned nodes, represented as the dashed red line at 2.21 cm in Fig. 17c. This identical arrival time is due to the same length of the myelin sheath and nodes of Ranvier. With the ephaptic coupling effect, the neural spike in each fiber bundle reaches its final node at different arrival times. The arrival time difference between the two fibers is 0.0202 ms for the same fiber bundle, represented as the solid black line at 2.21 cm in Fig. 17c. This arrival time difference is significant, approximately 26.5 times larger than the propagation without ephaptic coupling.

The neural spike in the first fiber bundle initially propagates faster than that in the second fiber bundle, as presented in Fig. 17b. This distinctive propagation difference is generated by the second myelin in *y* ∈ [0.13, 0.34] of the first fiber bundle, which is twice as lengthy as the second myelin in *y* ∈ [0.13, 0.23] of the second fiber bundle. The advanced propagation in the first fiber bundle constantly slows the lagged propagation via ephaptic coupling in the second fiber bundle. Ephaptic coupling dramatically changes the instantaneous conduction velocity in the two fiber bundles with misaligned nodes, compared to a single fiber bundle propagation, as confirmed in Fig. 17d. The last myelin in the second fiber bundle is twice as long as the last myelin in the first fiber bundle, making the total length the same.

Nevertheless, the modified initial propagation velocity by the various myelin lengths produces a different arrival time for both fiber bundles at each final node. If the fiber is propagated reversely from the last node of Ranvier, the neural spike in the second fiber bundle would always be faster than that of the first. The direction of the propagation determines which neural spikes propagates faster in the same two fiber bundles. However, in both directions, the synchronization between these two misaligned fiber bundles is almost impossible unless ephaptic coupling is negligible.

Moreover, this work considered a small delayed initiation in either the first or second fiber bundle of 0.01 ms for each fiber bundle. When the first fiber bundle is delayed by 0.01 ms, Fig. 18a demonstrates that the arrival time difference between the fiber bundles is 0.0147 ms. The arrival time difference is 0.0055 ms shorter than that by the simultaneous initiation. Only a fraction of the delay of the first fiber bundle of 0.01 ms is reflected in the arrival time difference. If the neural spike initiation in the second fiber bundle is delayed by 0.01 ms, the arrival time difference between the fiber bundles is 0.0246 ms, 0.0043 *ms* longer than that via the simultaneous initiation. Similar to the delay of the first fiber bundle, only a fraction of that of the second fiber bundle of 0.01 ms is reflected in the arrival time difference. Thus, the magnitude of the initiation delay is not linear to the arrival time difference in a misaligned fiber. This nonlinear phenomenon seems to be due to the dramatically changed conduction velocity owing to the complexity of ephaptic coupling between the nodes of Ranvier, as illustrated in Figs. 18b and 18c.

**Figure 18:**
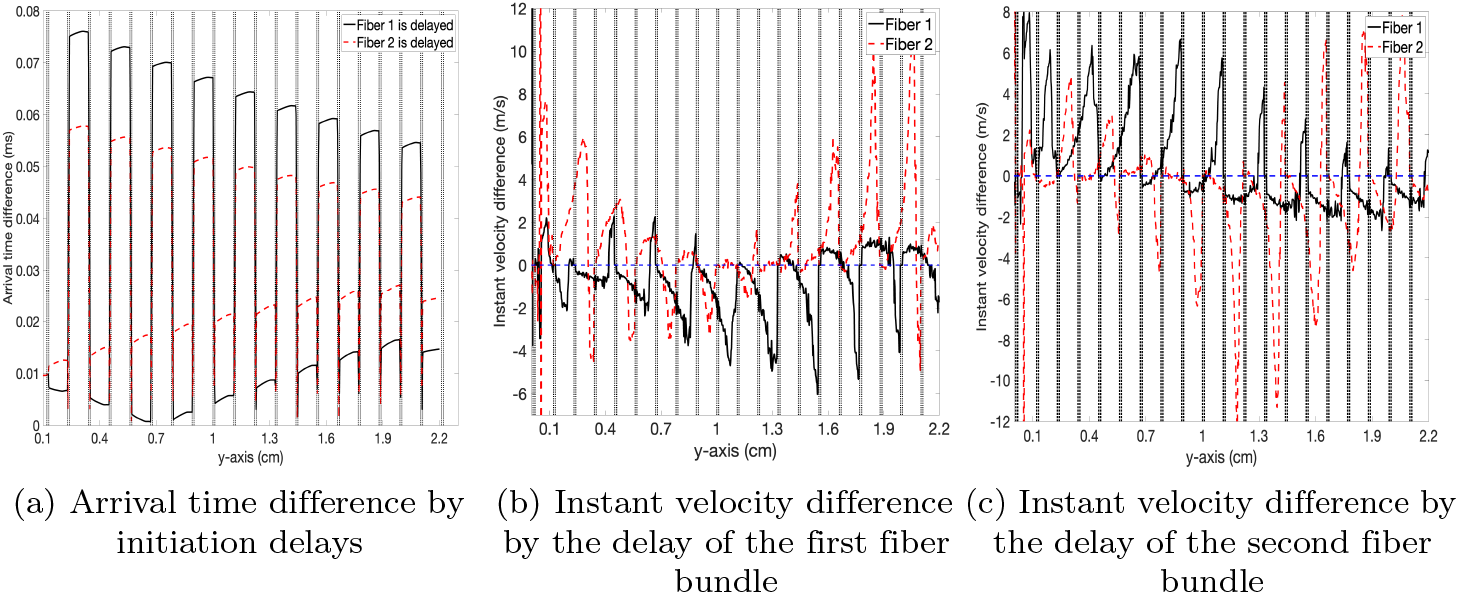
Arrival time and instant velocity of the misaligned fibers with a delay of 0.01 ms for either fiber bundle. (a) Arrival time difference by the delay of the first and second fiber bundle 0.01 *ms*. (b) Instant velocity changes of the two fiber bundles by the delay of the first fiber bundle 0.01 ms, compared to a single fiber bundle propagation. (c) Instant velocity changes in the two fiber bundles by the delay of the second fiber bundle 0.01 *ms*, compared to a single fiber bundle propagation where *ρ* = 1, *r*_*b*_ = 0.1 cm, and *T* = 24°*C*.

### E. Curvature effect: Two curved fiber bundles

The final case in the multidimensional bidomain model is the neural spike propagation in curved fiber bundles. This work considers two curved fiber bundles embedded in a rectangular 2D domain of (*x, y*) ∈ [0, 1] × [0, 1]. The curved fiber bundle is a circular shell with a constant radius *r*, centered at [0, 0], and the fiber width is significantly small. Thus, the fiber curvature is approximately constant. For convenience, the polar coordinate system (*r, θ*) is introduced to represent the fiber bundle geometry, where *θ* advances counter-clockwise. The first fiber bundle lies in *r* ∈ [0.81, 0.82] and *θ* ∈ [0.075, 1.388], whereas the second fiber bundle lies in *r* ∈ [0.83, 0.84] and *θ* ∈ [0.075, 1.388]. The node and myelin lengths are similar to the previous linear fiber bundles, approximately 0.2 ms and 0.01 ms, respectively. The initial excitation is assumed to start at the nodes of *θ* = [0.0561, 0.0686] in both fiber bundles.

The extracellular potential distribution of the curved fiber bundles is not linear, as presented in Fig. 19. A Poisson distribution with a few sources at the nodes along the curved fiber bundle generates a unique extracellular potential distribution. The distribution at T=0.6 ms is striking because the extracellular potential distribution affects the membrane potential in the backward part of the propagation in the fiber bundle. The nonlinear distribution of the extracellular potential deforms the propagation features of its own and neighboring fiber bundles’s. This distribution contrasts the extracellular potential of the linear fiber bundles, which is approximately linear and orthogonal to the membrane and boundary (Fig. 20).

**Figure 19:**
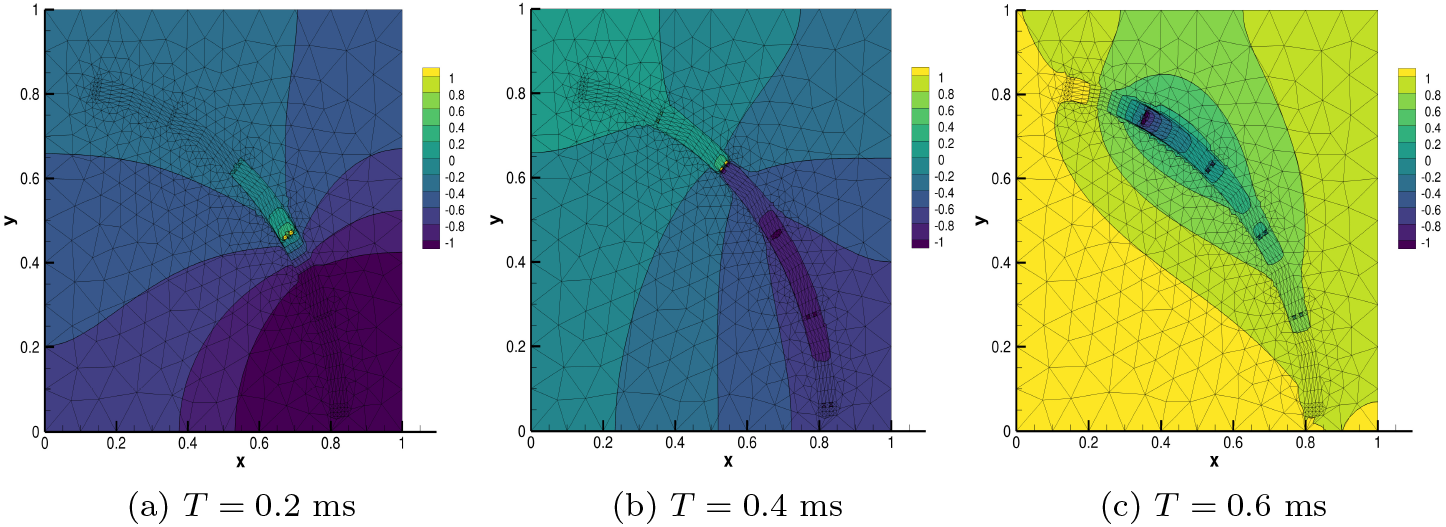
Extracellular potential distribution for the propagation in the domain embedding for two curved fiber bundles. (a) Extracellular potential distribution at (a) *T* = 0.2 ms, (b) *T* = 0.4 ms, (c) *T* = 0.6 ms. The neural spike initiates at *θ* ∈ [0.0561, 0.0686], where *ρ* = 1, *r*_*b*_ = 0.1 cm, and *T* = 24°*C*.

**Figure 20:**
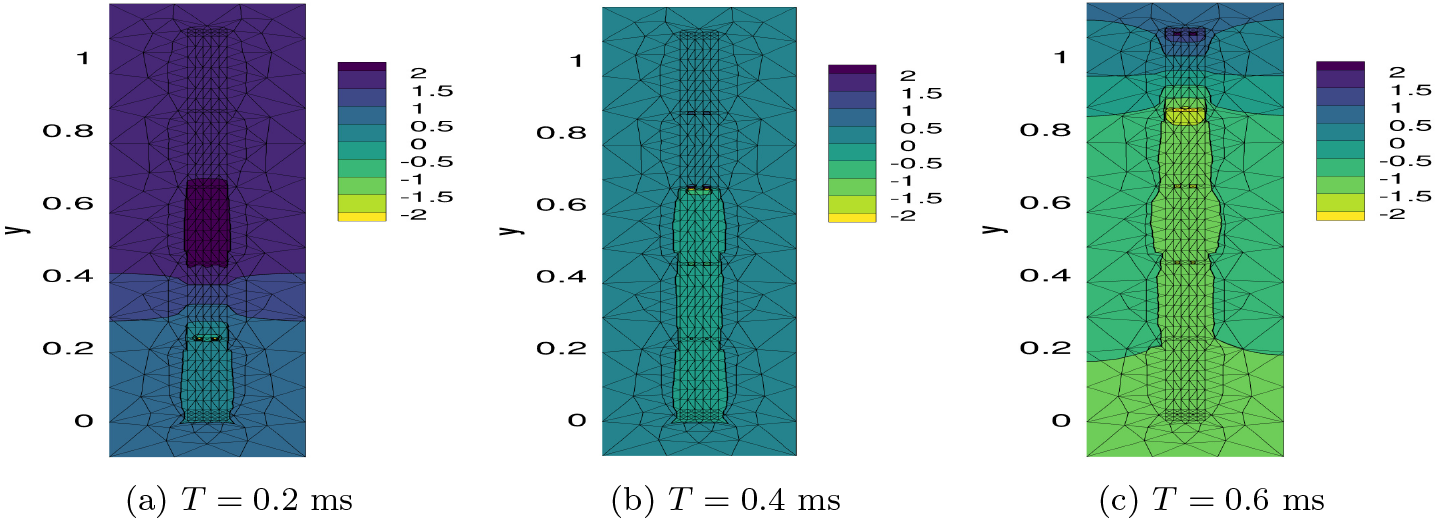
Extracellular potential distribution for the propagation in the domain embedding for two fiber bundles. (a) Extracellular potential distribution at (a) *T* = 0.2 ms, (b) *T* = 0.4 ms, (c) *T* = 0.6 ms. The neural spike initiates at *y* ∈ [0.01, 0.02], where *ρ* = 1, *r*_*b*_ = 0.1 cm, and *T* = 24°*C*.

Next, Fig. 21 demonstrates the arrival time and conduction velocity difference between the two fiber bundles, compared to the propagations in a single fiber bundle and the propagation without ephaptic coupling. In Fig. 21a, the single curved fiber bundle with ephaptic coupling (blue line) displays a similar conduction velocity along each fiber bundle. The slight difference in the arrival time for the same *θ* value is caused by the slight radius difference. The two fiber bundles with ephaptic coupling (black line) illustrate the decreased arrival time difference, indicating that the second fiber with a slightly larger radius has a higher conduction velocity. This phenomenon is coincident with ephaptic coupling for linear fiber bundles. In addition, Fig. 21b displays the instantaneous conduction velocity difference between the two fiber bundles. This figure confirms that the second fiber bundle speeds up during propagation along curved fiber bundles. Therefore, the synchronized neural spikes along curved fiber bundles maintain synchronization at the first nodes, although the total length of the fiber bundles differs. Detailed features of neural propagation in curved fiber bundles are beyond the scope of this paper and will be studied in future work.

**Figure 21:**
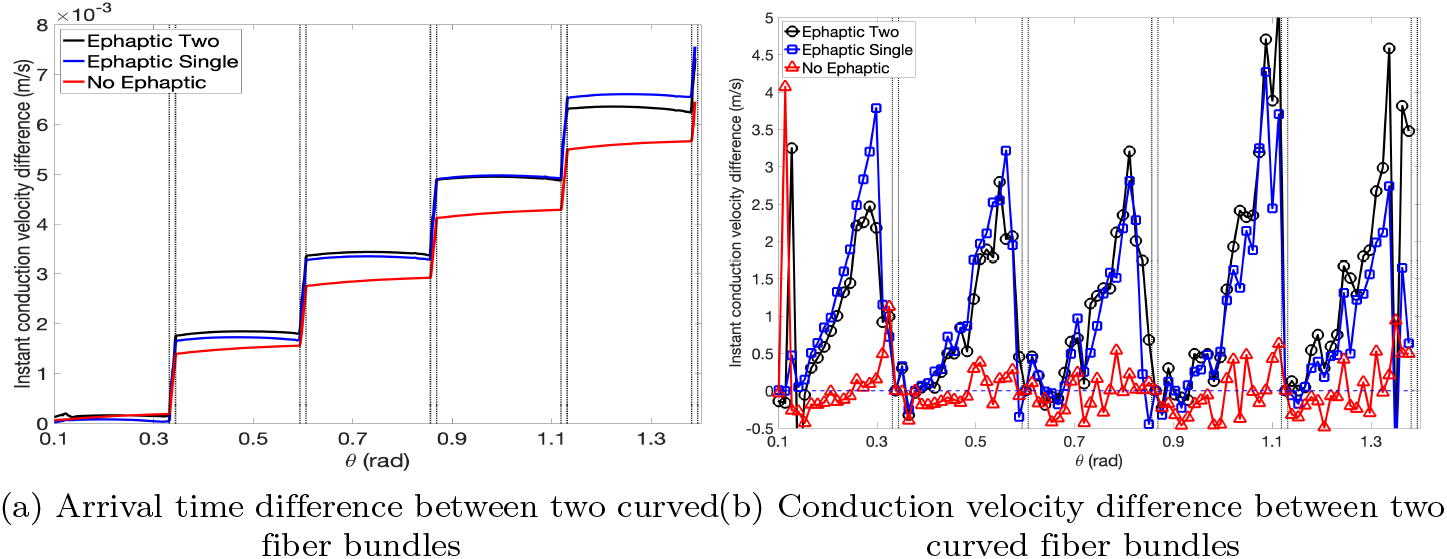
Arrival time and conduction velocity difference between the two curved fibers. Two fiber bundles with ephaptic coupling (black), a single fiber bundle with ephaptic coupling (blue), and propagation without ephaptic coupling (red). (a) Arrival time difference in milliseconds, and (b) conduction velocity difference in meters per second, where *ρ* = 1, *r*_*b*_ = 0.1 cm, and *T* = 24°*C*.

## 4 Discussion

This paper proposes a multidimensional bidomain model for neural spike propagation to study the ephaptic coupling effects in a multidimensional complex domain. In contrast to the 1D differential equations for neural spike propagation studies, the proposed model is expressed as partial differential equations for the general domain of multi-dimensional space. Orthonormal basis vectors (moving frames) are employed in the numerical scheme to tackle the two separate domains for the membrane and extracellular potential. Computational simulations in 2D domains for the known results are provided to validate the proposed model and confirm the known results in a 1D domain. Moreover, additional simulations are provided on three new cases which are only possible in a multidimensional model; interference by opposite-traveling neural spikes, propagation in misaligned neural fiber bundles, and curved fiber bundles.

The proposed model and corresponding numerical scheme have the following limitations. First, the ratio between the width and length of a single neural fiber is significantly large, such that the simulation of a single fiber is extremely expensive or impossible, due to numerous elements representing the geometry. As a solution, the proposed model instead fits a fiber bundle with a width of more than about 0.01 *cm*, which contains tens of thousands of identical neural fibers, although the strength of the extracellular potential should be adjusted accordingly. Second, the computational cost of solving the proposed model is relatively high, even in the domain of a fiber bundle with a reasonable number of elements. This high computational cost is caused by the numerical stiffness in (a) the ion channel reaction function, (b) the Laplacian operator with strong anisotropy, and (c) the repetitive anisotropic Poisson solver in the bidomain equation. The two fiber bundles with a length of 1.1 cm with approximately 500 triangular elements take about 50 hours for up to 1 ms at a workstation with a single central processing unit (CPU). A parallel CPU computation reduces the computational time inversely proportional to the number of CPUs. Further model reduction or 3D–1D modeling seem necessary to enhance computational efficiency, possibly for full 3D simulations of real fiber bundles of the human brain.

## Acknowledgments

The research of the first author is supported by the National Research Foundation of Korea (NRF-2021R1A2C109297811). The research of the second author is supported by JSPS Grant-in-Aid for Scientific Research (24K06852), JST CREST (JPMJCR1914), and Keio University (Fukuzawa Memorial Fund).

## References

1. C. Anastassiou and C. Koch. Ephaptic coupling to endogenous electric field activity: why bother? Curr. Opin. Neurobiol., 31:95–103, 2015.

2. C. A. Anastassiou, R. Perin, H. Markram, and C. Koch. Ephaptic coupling of cortical neurons. Nat. Neurosci., 14, 2011.

3. A. Arvanitaki. Effects evoked in an axon by the activity of a contiguous one. J. Neurophysiol., 5:89–108, 1942.

4. U. Ascher, S. Ruuth, and R. Spiteri. Implicit-explicit runge-kutta methods for time-dependent partial differential equations. Appl. Numer. Math., 25:151–167, 1997.

5. S. Binczak, J. C. Eilbeck, and A. C. Scott. Ephaptic coupling of myelinated nerve fibres. Physica D, 148:159–174, 2001.

6. G. Bizsáki, C. A. Anastassiou, and C. Koch. The origin of extracellular fields and currents - eeg, ecog, lfp and spikes. Nat. Rev. Neurosci., 13:407–420, 2012.

7. H. Bokil, N. Laaris, K. Blinder, M. Ennis, and A. Keller. Ephaptic interactions in the mammalian olfactory system. J. Neurosci., 21:1–5, 2001.

8. M. H. Brill, S. G. Waxman, J. W. Moore, and R. W. Joyner. Conduction velocity and spike configuration in myelinated fibres: computed dependence on internode distance. J. Neurol. Neurosurg. Psychiatry, 40(8):769–774, 1977.

9. M. Capllonch-Juan and F. SepulvedaI. Modelling the effects of ephaptic coupling on selectivity and response patterns during artificial stimulation of peripheral nerves. PLOS Comput. Biol., 16(6):1007826, 2020.

10. E. Cartan. Geometry of Riemannian Spaces. Math. Sci. Press, 2001.

11. E. Cartan. Riemannian Geometry in an Orthogonal Frame. World Scientific Pub. Co. Inc., 2002.

12. C.-C. Chiang, R. S. Shivacharan, X. Wei, L. E. Gonzalez-Reyes, and D. M. Durand. Slow periodic activity in the longitudinal hippocampal slice can self-propagate non-synaptically by a mechanism consistent with ephaptic coupling. J. Physiol., 591(1), 2018.

13. S. Chun. Method of moving frames to solve conservation laws on curved surfaces. J. Sci. Comput., 53(2):268–294, 2012.

14. S. Chun. Method of moving frames to solve (an)isotropic diffusion equations on curved surfaces. J. Sci. Comput., 59(3):626–666, 2013.

15. S. Chun. Method of moving frames to solve the time-dependent Maxwell’s equations on anisotropic curved surfaces: Applications to invisible cloak and ELF propagation. J. Compt. Phys., 340:85–104, 2017.

16. S. Chun and C. Eskilsson. Method of moving frames to solve the shallow water equations on arbitrary rotating curved surfaces. J. Compt. Phys., 333:1–23, 2017.

17. S. Chun and J. Jung. Method of time map spatializing the dynamics of biological electric signal propagation and its applications to quick multidimensional simulation and deformed conductivity reconstruction. submitted., 2024.

18. B. Frankenhaeuser and A. F. Huxley. The action potential in the myelinated nerve fibre of xenopus laevis as computed on the basis of voltage clamp data. J. Physiol., 171:302–315, 1964.

19. L. Goldman and J. S. Albus. Computation of impulse conduction in myelinated fibers; theoretical basis of the velocity-diameter relation. Biophys. J., 8(5):596–607, 1968.

20. J. E. Hall and M. E. Hall. Guyton and Hall Textbook of Medical Physiology. Elsevier, 11 edition, 2021.

21. P. E. Hand and B. E. Griffith. Adaptive multiscale model for simulating cardiac conduction. PNAS, 107(33):14603–14608, 2010.

22. A. L. Hodgkin and A. F. Huxley. Currents carried by sodium and potassium ions through the membrane of the giant axon of loligo. J. Physio., 116(4):449–72, 1952.

23. J. D. Jackson. Classical Electrodynamics. Wiley, third edition, 2021.

24. C. Jerez-Hanckes, I. Pettersson, and V. Rybalko. Derivation of cable equation by multiscale analysis for a model of myelinated axons. Discrete and Continuous Dynamical Systems - B, 25(3):815–839, 2020.

25. S. M. Kandel. The electrical bidomain model: A review. Sch. Acad. J. Biosci., 3(7):633–639, 2015.

26. B. Katz and O. H. Schmitt. Electric interaction between two adjacent nerve fibres. J. Physiol., 97:471–488, 1940.

27. R. W. Lewis, P. Nithiarasu, and K. N. Seetharamu. Fundamentals for the Finite Element Method for Heat and Fluid Flow. Wiley, 2004.

28. J. Lin and J. P. Keener. Modeling electrical activity of myocardial cells incorporating the effects of ephaptic coupling. PNAS, 107(49):20935–20940, 2010.

29. V. S. Markin. Electric interaction of parallel non-myelinated nerve fibers. 1. change in excitability of the adjacent fibre. Biofizika, 15(1):122–133, 1970.

30. V. S. Markin. Electric interaction of parallel non-myelinated nerve fibers. 2. shared conduction of impulses. Biofizika, 15(4):681–9, 1970.

31. V. S. Markin. Electric interaction of parallel non-myelinated nerve fibers. 3. interaction in nerve trunk. Biofizika, 18(2):314–321, 1973.

32. V. S. Markin. Electric interaction of parallel non-myelinated nerve fibers. 4. the role of anatomic non uniformities of the nerve trunk. Biofizika, 18(3):512–518, 1973.

33. A. S. Marrazzi and R. Lorente. Interaction of neighboring fibres in myelinated nerve. J. Neurophysiol., 7(2):83–101, 1944.

34. U. Mitzdorf. Current source-density method and application in cat cerebral cortex: investigation of evoked potentials and eeg phenomena. Physiol. Rev., 65(1):37–100, 1985.

35. Y. Mori, G. I. Fishman, and C. S. Peskin. Ephaptic conduction in a cardiac strand model with 3D electrodiffusion. PNAS, 105(17):6463–6468, 2008.

36. D. Moxey, C. D. Cantwell, Y. Bao, A. Cassinelli, G. Castiglioni, S. Chun, E. Juda, E. Kazemi, K. Lackhove, J. Marcon, G. Mengaldo, D. Serson, M. Turner, H. Xu, J. Peiró, R. M. Kirby, and S. J. Sherwin. Nektar++: enhancing the capability and application of high-fidelity spectral/hp element methods. Compt. Phys. Comm., 249:107110, 2020.

37. C. Nicholson and J. A. Freeman. Theory of current source-density analysis and determination of conductivity tensor for anuran cerebellum. J. Neurophys., 38:356–368, 1975.

38. V. K. Nielsen. Pathophysiology of hemifacial spasm: I. ephaptic transmission and ectopic excitation. Neurol., 34(4):418–426, 1984.

39. I. Pettersson, A. Rybalko, and V. Rybalko. Bidomain model for axon bundles with random geometry. Gas dynamics with applications in industry and life sciences, 2022.

40. W. Pitts. Investigations on synaptic transmission. Cyber. Trans. 9th Conf., pages 159–162, 1952.

41. A. J. Pullan, M. L. Buist, and L. K. Cheng. Mathematically modelling the electrical activity of the heart. World Scientific, 2005.

42. M. Rasminsky. Ephaptic transmission between single nerve fibres in the spinal roots of dystrophic mice. J. Physiol., 350:151–169, 1980.

43. S. Reutskiy, E. Rossoni, and B. Tirozzi. Conduction in bundles of demyelinated nerve fibers: computer simulation. Biol. Cybern., 89:439–448, 2003.

44. B. J. Roth. CElectrical conductivity values used with the bidomain model of cardiac tissue. IEEE Trans. Bio. Eng., 44(4):326–328, 1997.

45. B. J. Roth and D. L. Beaudoin. Approximate analytical solutions of the bidomain equations for electrical stimulation of cardiac tissue with curving fibers. Phys. Rev. E, 67:051925, 2003.

46. S. Rush and H. Larsen. A practical algorithm for solving dynamic membrane equations. IEEE Trans. Bio. Eng., 25(4):389–392, 1978.

47. R. M. Sapolsky. Determined: A Science of Life without Free Will. Penguin Press, 2023.

48. H. Schmidt, G. Hahn, G. Deco, and T. R. Knösche. Ephaptic coupling in whiter matter fibre bundles modulates axonal transmission delays. PLOS Comput. Biol., 17(2):e1007004, 2021.

49. H. Schmidt and T. R. Knösche. Action potential propagation and synchronisation in myelinated axons. PLOS Comput. Biol., 15(10):e1007004, 2019.

50. H. Schmidt and T. R. Knösche. Modelling the effect of ephaptic coupling on spike propagation in peripheral nerve fibres. Biol. Cyber., 116:461–473, 2022.

51. O. H. Schmitt. Information processing in the nervous system. Proceedings of a symposium held at the State University of New York at Buffalo, 21st-24th October, 1968.

52. H. Sheheitli and V. K. Jirsa. A mathematical model of ephaptic interactions in neuronal fiber pathways: Could there be more than transmission along the tracts? Netw. Neurosci., 4(3):595–610, 2020.

53. I. Tasaki. Nervous Trasmission. Thomas, 1953.

54. C. Taylor and F. Dudek. Synchronous neural afterdischarges in rat hippocampal slices without active chemical synapses. Science, 218(4574):810–812, 1982.

55. L. Tung. A Bi-Domain Model for Describing Ischemic Myocardial D-C Potentials. Phd thesis, MIT, 1978.

56. H. Versteeg and W. Malalasekra. An Introduction to Computational Fluid Dynamics. Pearson, 2007.

57. S. G. Waxman. Conduction in myelinated, unmyelinated, and demyelinated fibers. Arch Neurol., 34(10):585–9, 1977.

58. S. G. Waxman and M. H. Brill. Conduction through demyelinated plaques in multiple sclerosis: computer simulations of facilitation by short internodes. J. Neurol., 41:408–416, 1978.

